# Gasdermin E couples mitochondrial stress to STING-driven neuronal pyroptosis during Chandipura virus encephalitis

**DOI:** 10.64898/2026.09.25.754282

**Authors:** Nripa Kumari, Sourav Dutta, Akasmika Priyadarsini, Joyanta Biswas, Shubhangi Roy, Suman Samanta, Semanti Chakraborty, Jyotirmoyee Roy, Anirban Basu, Debanjan Mukhopadhyay, Abhik Saha, Piyali Mukherjee

## Abstract

Neurotropic RNA viruses are major causes of fatal encephalitis worldwide, yet how infected neurons transition from antiviral defense to inflammatory cell death is not well characterized. Chandipura virus (CHPV), an emerging neurotropic rhabdovirus, causes acute, rapidly progressive encephalitis with high case fatality in children, but the mechanisms underlying its neuropathogenesis remain poorly defined. Here, we demonstrate that CHPV suppresses canonical RNA virus sensing early but subsequently switches to a mitochondria-driven innate immune program that culminates in inflammatory cell death. Early infection of neuronal cells with CHPV was marked by reduced levels of the mitochondrial antiviral adaptor protein, MAVS and attenuation of type I and III interferon responses. As infection progressed, mitochondrial dysfunction promoted accumulation of mtROS, mitochondrial accumulation of cleaved GSDME and cytosolic mtDNA release, triggering STING activation, which coincided with robust neuroinflammation and pyroptotic cell death. Pharmacological inhibition or genetic silencing of STING markedly attenuated inflammatory signaling, prevented pyroptotic membrane rupture and protected neurons from cell death without significantly affecting viral replication. In contrast, GSDME depletion reduced both viral replication and neuronal death. Notably, GSDME depletion markedly attenuated STING phosphorylation, while STING depletion also reduced GSDME activation, revealing functional coupling between these pathways during CHPV-induced neuronal injury. Collectively, our findings identify a mitochondria-GSDME-STING axis linking early immune evasion to neuroinflammation during CHPV infection, revealing a previously unrecognized mechanism of inflammatory neuronal death in viral encephalitis and highlighting STING as a potential therapeutic target in certain CNS viral infections.

**Author summary:** Members of the *Rhabdoviridae* family, including Rabies virus and CHPV, can cause acute encephalitis with high case-fatality rates. Despite their clinical burden, effective therapeutic options for CHPV-associated encephalitis remain limited. Recent CHPV outbreaks in India, particularly among children, highlight the need to better understand the host responses that contribute to disease progression. Here, we show that CHPV infection induces a temporally distinct antiviral and inflammatory response, characterized by early suppression of mitochondrial antiviral signaling followed by mitochondrial dysfunction, mtDNA release, and robust STING activation. This transition is associated with mitochondrial accumulation and cleavage of GSDME. GSDME depletion reduced STING activation, while STING depletion reciprocally reduced GSDME cleavage, revealing a functional interplay between these pathways. Although caspase-1 was activated, we did not detect convincing GSDMD cleavage, suggesting that CHPV-induced neuronal inflammatory cell death is not fully explained by the canonical inflammasome pathway. Cell-type-specific responses in the infected brain further highlight the complexity of CHPV-associated neuroinflammation. Together, these findings identify the GSDME-STING axis as a potential therapeutic target to limit pathological neuroinflammation and neuronal injury during CHPV encephalitis.

## Introduction

The central nervous system (CNS) is a tightly regulated, immune-privileged environment, safeguarded by the blood-brain barrier (BBB) and limited immune surveillance to preserve tissue integrity [1]. Despite these constraints, several neurotropic RNA viruses, including members of the *Rhabdoviridae, Flaviviridae, Paramyxoviridae, Picornaviridae* and *Coronaviridae* families, have evolved diverse strategies to access the CNS, including hematogenous dissemination across a compromised BBB, “Trojan horse” entry via infected immune cells, and retrograde axonal transport through peripheral nerves [2–5]. Infection of the CNS by these viruses frequently culminates in viral encephalitis, a severe and often fatal condition characterized by widespread neuroinflammation, cellular dysfunction and neurological impairment. Classical encephalitic viruses such as Japanese encephalitis virus (JEV) and West Nile virus (WNV) remain major global threats of viral encephalitis [6–8]. Chandipura virus (CHPV), a neurotropic pathogen of the *Rhabdoviridae* family, is an emerging concern, particularly in India, where recurrent outbreaks with substantial pediatric mortality have prompted renewed national efforts to understand its transmission and pathogenesis (https://www.pib.gov.in/PressReleasePage.aspx?PRID=2294690&reg=48&lang=2). The disease can progress rapidly from an acute febrile illness to severe encephalitis and death, particularly in children, while no specific antiviral therapy is currently available [9, 10]. This combination of rapid disease progression, high mortality and limited therapeutic options highlights the need to understand how CHPV interacts with host pathways to drive neurological injury.

At the cellular level, neurons and other CNS-resident cells are now recognized as active participants in innate immune responses, capable of sensing viral infection and initiating inflammatory signaling [11, 12]. However, the restricted regenerative capacity of neural tissue renders the CNS particularly vulnerable to immune-mediated damage, where even transient dysregulation of inflammatory responses can result in irreversible functional loss and long-term neurological sequelae. Consequently, the antiviral signaling pathways activated within neurons must be precisely regulated to ensure effective pathogen clearance while minimizing tissue damage. The first line of defense against RNA viruses within the cells is mediated by cytosolic sensors of the retinoic acid-inducible gene-I (RIG-I)-like receptor (RLR) family [13]. Upon recognition of viral RNA, RIG-I engages the mitochondrial adaptor protein MAVS, leading to the assembly of a multiprotein signalosome comprising TRAF2, TRAF3, TRAF5 and TRAF6 along with the TANK-binding kinase 1 (TBK1), ultimately driving IRF3/7 phosphorylation and NF-κB-dependent transcription of type I/III interferons (IFNs) and pro-inflammatory cytokines [14, 15]. While this pathway is essential for antiviral defense, neurotropic RNA viruses have evolved mechanisms to attenuate MAVS signaling and delay IFN responses for establishing successful infection [16, 17]. For example, JEV interferes with MAVS-dependent signaling through its nonstructural proteins and host ubiquitin-mediated pathways [18, 19], whereas WNV dampens downstream signaling outputs without directly targeting MAVS for cleavage, thereby modulating rather than abolishing antiviral responses [20, 21]. In contrast, for CHPV, mechanistic insights into MAVS regulation remain limited despite evidence of potent IFN antagonism [22–24], representing a critical gap in understanding host-virus interactions in emerging encephalitic infections. Additionally, whether suppression of the canonical RLR-MAVS pathway is subsequently compensated by alternative innate immune sensing mechanisms during neurotropic RNA virus infection, especially during CHPV infection, remains largely unknown.

Beyond canonical RLR-MAVS signaling, accumulating evidence indicates that RNA virus infections can engage alternative innate immune pathways, including the cyclic GMP-AMP synthase (cGAS)-STING axis [25, 26]. Although traditionally associated with DNA sensing, it is increasingly recognized that STING can be activated during RNA virus infection through the release of intracellular DAMPs like mitochondrial DNA (mtDNA) under conditions of cellular stress [27–29]. Notably, STING signaling has been linked not only to IFN responses but also to inflammatory cell death pathways, particularly pyroptosis mediated by gasdermin (GSDM) family proteins [30–31]. GSDMs function as key executors of membrane permeabilization, with Gasdermin D (GSDMD) classically associated with inflammasome activation, while Gasdermin E (GSDME) links apoptotic and pyroptotic signaling through caspase-3-dependent mechanisms [32]. In viral infections, STING is widely recognized as a host-protective antiviral pathway that promotes immune activation and viral clearance, as demonstrated in neurotropic viruses such as JEV [33]. However, whether such inflammatory cell death is uniformly beneficial within the CNS remains unclear. Given the limited regenerative capacity and functional sensitivity of neural tissue, excessive or deregulated GSDM activation may exacerbate neuronal damage and contribute to disease pathogenesis. Furthermore, while inflammasome activation has been described in response to several RNA viruses [34], the upstream signals that couple mitochondrial perturbation to STING activation and subsequent GSDM-mediated membrane permeabilization is not defined, and the possibility of temporally distinct engagement of different GSDM family members has not been explored. Importantly, in the context of CNS infection, such a shift from antiviral signaling to inflammatory cell death may represent a key determinant of neuropathology.

In this study, we demonstrate that CHPV elicits a temporally orchestrated innate immune response in neurons, characterized by an early suppression of MAVS-dependent antiviral signaling followed by activation of a mitochondria-derived inflammatory program. During the early phase of infection, downregulation of MAVS is accompanied by attenuated type I and III IFN responses, which probably facilitates viral establishment. As infection progresses, mitochondrial dysfunction promotes the selective cleavage of GSDME at the mitochondria, cytosolic release of mtDNA and STING activation that coincides with pyroptotic neuronal death. Importantly, this signaling axis was validated in a mouse model of CHPV encephalitis, highlighting its physiological relevance in vivo. In contrast to its well-established antiviral role, our findings reveal that STING activation in CHPV-infected neurons does not significantly restrict viral replication but instead drives inflammatory neuronal injury. Thus, our study reveals a temporal transition from an early, deregulated antiviral response to a mitochondria-associated inflammatory cell death program involving STING and GSDME and highlight STING as a potential target for host-directed therapeutic intervention during viral encephalitis.

## Results

### CHPV exhibits selective neuronal tropism accompanied by progressive viral replication and neurodegeneration both in vivo and in vitro

Neurotropic RNA viruses are characterized by their ability to cross the BBB and access the CNS, where they infect resident cell populations and contribute to neuroinflammation and altered brain function [35]. However, whether virus-induced neuronal death represents a host-driven antiviral strategy to limit viral replication or a virus-mediated mechanism to facilitate dissemination within the CNS remains unresolved. CHPV is a highly neurotropic rhabdovirus that causes acute encephalitis with selective neuronal injury in children. Previous studies have established that CHPV productively infects neuronal cells and induces apoptosis through Fas/FADD-mediated signaling [36, 37], while parallel studies have demonstrated activation of inflammatory responses in microglia [38]. However, the molecular events that coordinate antiviral innate immune signaling with inflammatory responses and subsequent neuronal death remain poorly understood. Further, much of the experimental work on CHPV has relied on non-neuronal cell lines, including Vero (African green monkey kidney epithelial cells) and MEF (mouse embryonic fibroblast) cells [39, 22]. While these systems have been instrumental in understanding viral replication, they do not adequately model neuronal cell biology. To delineate the molecular mechanisms driving neuronal cell death during CHPV infection, we first examined the temporal dynamics of CHPV infection *in vivo*. Ten-day old BALB/c mice were challenged intraperitoneally with 3000 pfu of CHPV and monitored at days 1 (D1) and 2 (D2) post-infection (dpi) and at full symptom (FS) (**Fig. 1A**). CHPV-infected animals exhibited progressive weight loss, increasing neurological symptom scores and higher mortality at later stages of infection (**Fig. S1A-C**). Quantitative RT-PCR (qRT-PCR) analysis revealed a progressive increase in viral burden within the brain and spinal cord of the infected animals, indicating robust CNS invasion and productive viral replication during disease progression (**Fig. 1B-C**). To determine the pathological consequences of progressive CHPV neuroinfection, histopathological examination of brain sections was performed with hematoxylin and eosin (H&E) staining. At D1, mild neuropathological alterations were evident as compared to the control brains, which exhibited normal pathology (**Fig. 1D**). At D2, pronounced vacuolation and disruption of the brain parenchyma was observed and mice at the FS stage displayed severe encephalitic pathology accompanied by extensive vacuolation and marked tissue degeneration (**Fig. 1D**). Consistent with these observations, immunofluorescence analysis revealed reduction in NeuN immunoreactivity within the cortical region of the brain as the disease progressed (**Fig. 1E**). These changes coincided with increasing CHPV immunoreactivity peaking at D2, suggesting that progressive viral replication is associated with neuronal damage during CHPV infection. We also observed a progressive disruption of the hippocampal neuronal layers following CHPV infection (**Fig. 1F**). In control mice, the hippocampal formation displayed well-defined dentate gyrus (DG) and cornu ammonis (CA) neuronal layers with robust NeuN immunoreactivity (**Fig. 1F**). At D1, the overall organization of the hippocampus remained largely preserved without appreciable alterations. By D2, early changes became evident, but these alterations were markedly exacerbated at the FS stage, with pronounced disorganization of the DG and CA regions and diminished NeuN immunoreactivity indicative of widespread neurodegeneration (**Fig. 1F**). Further, Iba1 staining (indicative of microglial population) showed increased branched structure round the NeuN stained cells in the cerebral cortex of mouse during the early stages of CHPV infection as compared to control mice that could be due to increased activation/infiltration of microglial population (**Fig. 1G**).

**Figure 1.**
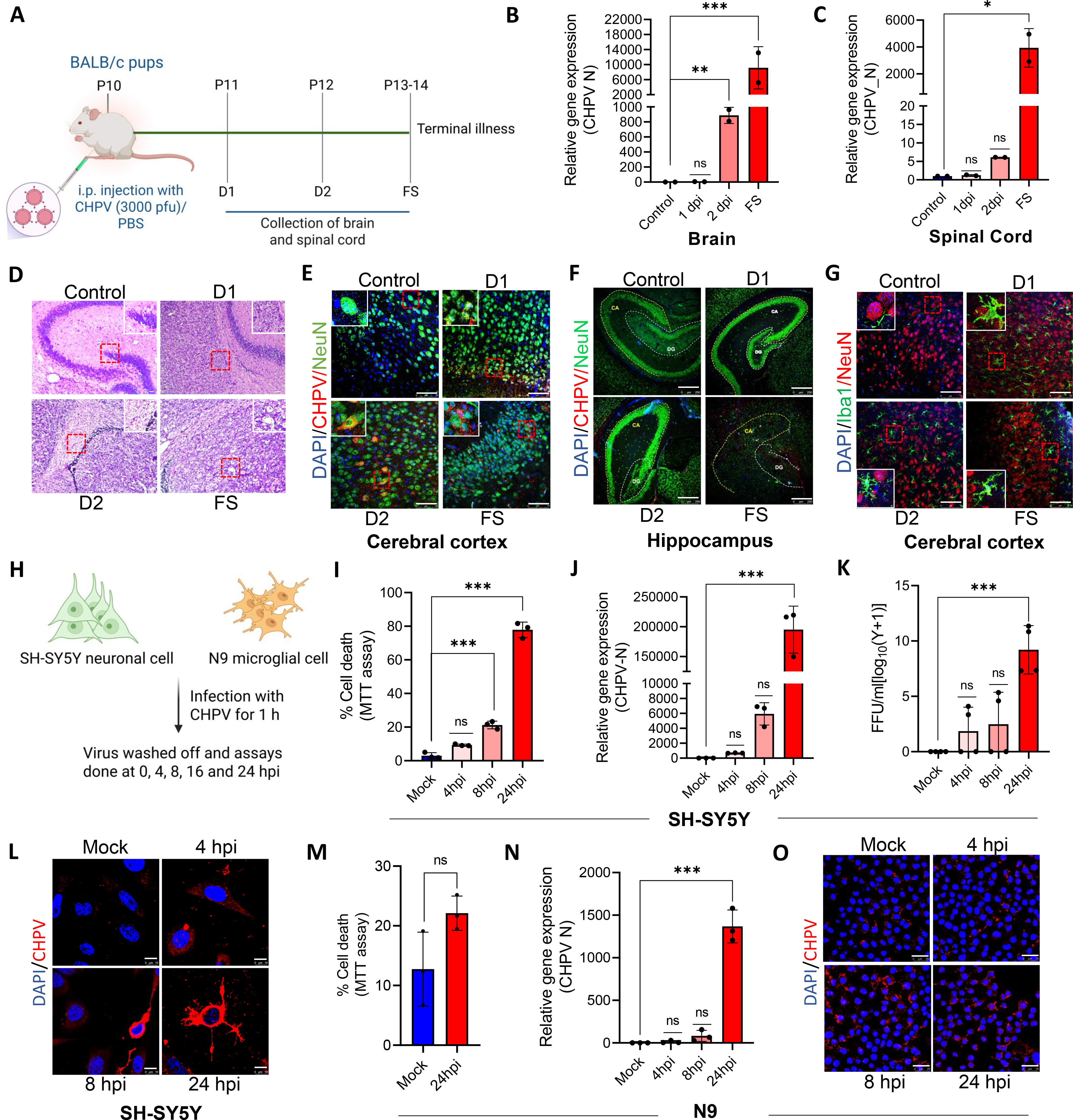
CHPV exhibits selective neuronal tropism accompanied by progressive viral replication and neurodegeneration. **(A)** Schematic outline of the experimental strategy for CHPV infection in BALB/c mice. **(B-C)** qRT-PCR analysis of relative gene expression of CHPV nucleocapsid (N) gene in mouse brain tissue (B) and mouse spinal cord tissue (C) at 1-day post-infection (dpi) (D1), 2-dpi (D2) and at full symptom (FS). (n=2 mice per group). **(D)** Haematoxylin and eosin staining of brain tissue of CHPV-infected mice at D1, D2 and FS and compared to control. (Scale bar = 10 µm). **(E)** Representative confocal images of brain tissue at D1, D2 and FS sections co-stained with CHPV N protein (red) and the neuronal marker NeuN (green) alongwith DAPI (Blue). (Scale bar = 75 µm). **(F)** Representative confocal images of brain tissue sections at D1, D2 and FS showing the hippocampal region co-stained with CHPV N protein (red) and NeuN (green) with alongwith DAPI (Blue). (Scale bar = 250 µm). **(G)** Representative confocal images of brain tissue sections at D1, D2 and FS co-stained with the microglial marker Iba1 (green) and the neuronal marker NeuN (red) alongwith DAPI (Blue). (Scale bar = 75 µm). **(H)** Schematic outline of the experimental strategy of CHPV infection in the neuronal SH-SY5Y cells and the microglial N9 cells. **(I)** Percentage of cell death in SH-SY5Y cells post CHPV infection (MOI-0.1) at 4, 8 and 24 hpi, measured by MTT assay and compared with mock-infected cells. **(J)** qRT-PCR analysis of the relative gene expression of the CHPV N gene in SH-SY5Y cells post CHPV infection (MOI-0.1) at 4, 8 and 24 hpi, compared with mock-infected cells. **(K)** Extracellular viral load measured by TCID assay from the supernatant of infected SH-SY5Y cells post-CHPV infection (MOI-0.1) at 4, 8 and 24 hpi compared with mock-infected cells. **(L)** Representative confocal images of SH-SY5Y cells stained with CHPV N protein (red) and DAPI (blue). (Scale bar = 10 µm). **(M)** Percentage of cell death in N9 microglial cells post-CHPV infection (MOI-0.1) at 24 hpi measured by MTT assay. Statistical significance was determined by student’s t-test. (**N)** qRT-PCR analysis of the relative gene expression of CHPV N gene in N9 cells post-CHPV infection (MOI-0.1) at 4, 8 and 24 hpi compared with mock-infected cells. **(O)** Representative confocal images of N9 cells stained against CHPV N protein (red) and DAPI (blue). (Scale bar = 25 µm). Data are presented as mean ± SD. Statistical significance was determined by one-way ANOVA with post hoc Tukey’s multiple comparison test (**P < 0.01; ***P < 0.001; n=3 biological replicates).

To further investigate the temporal dynamics of CHPV infection in distinct CNS-resident cell populations, human neuronal SH-SY5Y cells and microglial N9 cells were infected with CHPV at 0.1 MOI upto 24 h (**Fig. 1H**). CHPV infection caused a progressive decline in neuronal viability, as assessed by MTT assay at 4, 8 and 24 hpi, concomitant with a time-dependent increase in viral load (**Fig. 1I–J**). Consistently, viral titers quantified as focus-forming units (FFU) showed a significant increase at 24 hpi (**Fig. 1K**), indicating robust production of infectious viral progeny. This was accompanied by enhanced viral spread within neuronal processes, as evidenced by increased CHPV staining along neurites as the infection progressed (**Fig. 1L**). In contrast, infection of N9 microglial cells under similar conditions did not result in a significant loss of cell viability (**Fig. 1M**). Despite a substantial increase in viral load at 24 hpi in N9 cells, CHPV replication remains ∼80 folds lower as compared to the SH-SY5Y cells (**Fig. 1N-O**), suggesting that neuronal cells are more susceptible to CHPV-induced cytotoxicity.

### Single-cell transcriptomic analysis of CHPV-infected mouse brain reveals cell-type-specific antiviral and inflammatory responses

To define the transcriptional landscape of CNS cell populations during CHPV infection, we performed single-cell RNA sequencing (scRNA seq) of whole brains from CHPV-infected mice at the FS stage and age-matched control mice (n=2 mice per group). Uniform Manifold Approximation and Projection (UMAP) of the integrated scRNA seq dataset revealed a dramatic change in the transcriptional response in both the neuronal and non-neuronal populations between uninfected and infected mice indicating a spatial reorganization of the brain microenvironment. (**Fig. 2A-B**). Comparison of the cellular composition between individual samples revealed differences in the relative representation of several cell populations including cerebellar granule neurons (CGN), olfactory bulb (OB) GABAergic neurons, striatal medium spiny neurons, cortical neurons, astrocytes, microglia, oligodendrocyte precursor cells (OPCs), oligodendrocytes and vascular and leptomeningeal cells (**Fig. 2C**). We next examined cell-type-specific antiviral and innate immune transcriptional responses following CHPV infection. Microglia exhibited robust induction of IFN-stimulated and antiviral genes, including *Isg15, Ifit1, Ifit3, Ifih1, Rsad2, Oasl2, Ddx60 and Irf7* (**Fig. 2D**). In contrast, astrocytes showed relatively limited changes in the expression of these antiviral genes. Notably, cerebellar granule neurons (CGN) displayed a distinct antiviral transcriptional response characterized by significant upregulation of *Stat1, Stat2, Stat3, Oasl2, Ddx60* and *Irgm1/2* (**Fig. 2D**). This response was markedly different from that observed in several other neuronal populations, including olfactory bulb GABAergic neurons (OB-in and OB-out), in which several antiviral genes, including Oasl2 and Ddx60, were significantly downregulated, while most other neuronal populations showed relatively limited changes (**Fig. 2D**).

**Figure 2.**
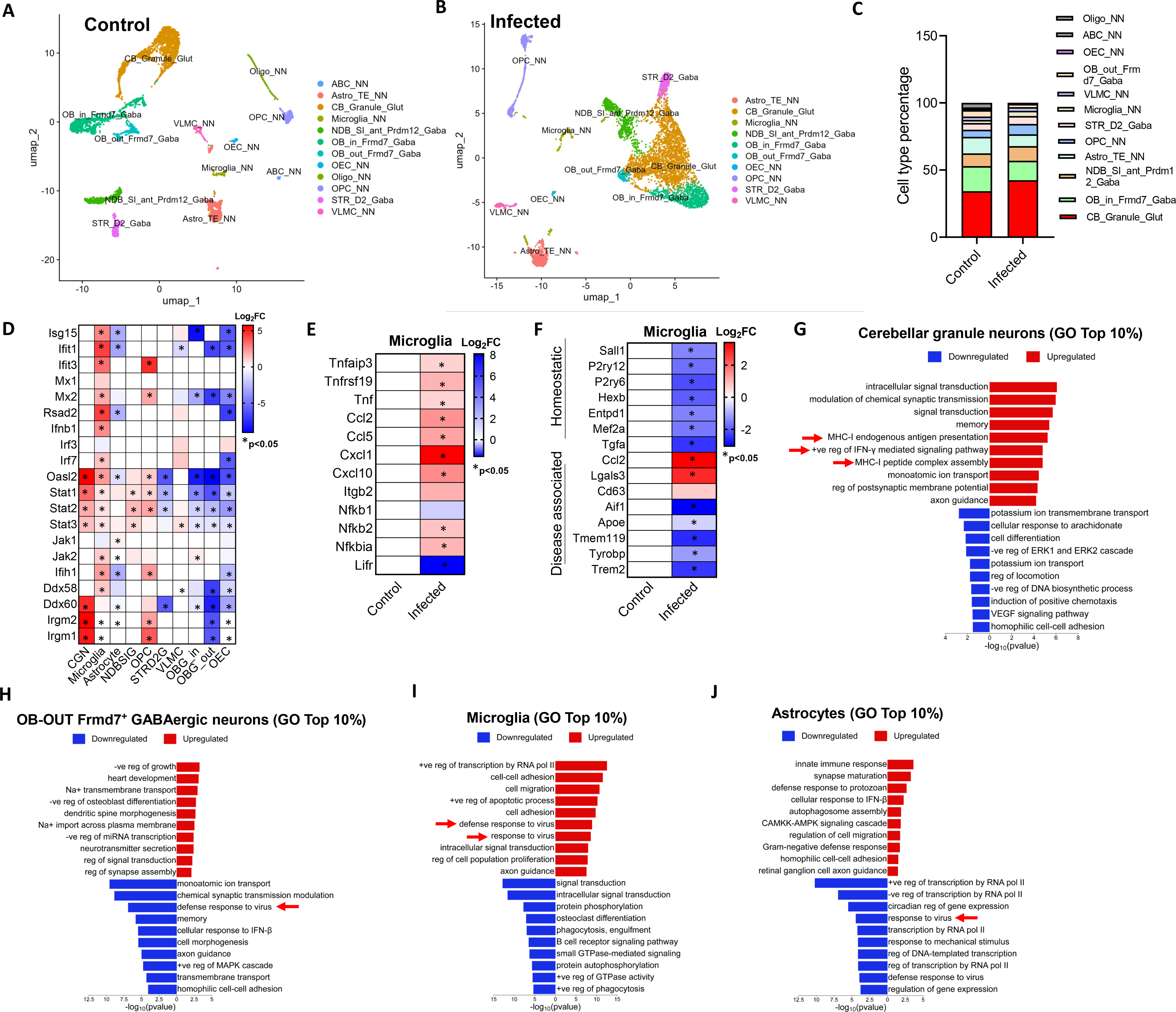
Single-cell transcriptomic analysis of CHPV-infected mouse brain reveals cell-type-specific antiviral and inflammatory responses. (A-B) UMAP visualization of single-cell transcriptomic profiles from control and CHPV-infected mouse brains (FS), showing the major neuronal and non-neuronal cell populations identified based on transcriptional signatures. (n=2 mice per group). (C) Cell-type-specific feature plots showing the distribution of selected cells in control vs CHPV-infected brain samples. (D) Heatmap showing relative mRNA expression of differentially expressed genes of the interferon pathway across the identified clusters. (E) Heatmap showing relative mRNA expression of differentially expressed genes of the inflammatory pathway in the microglial cluster. (F) Heatmap showing relative mRNA expression of differentially expressed genes depicting homeostatic vs disease-associated genes in the identified microglial cluster. (G-J) Gene Ontology (GO) enrichment analysis (top 10%) highlighting infection-associated changes in the identified cellular clusters following CHPV infection. Enrichment is represented as −log_10_(*P* value), with magenta and green indicating upregulated and downregulated biological processes, respectively in the indicated cellular clusters.

In addition to the antiviral response, CHPV infection induced a prominent inflammatory transcriptional program in microglia, characterized by significant increase in the expression of inflammatory markers like *Ccl2, Ccl5, Cxcl1, Cxcl10, Lifr* and *Nfkbia* (**Fig. 2E**). We therefore examined genes associated with both homeostatic microglial identity and disease-associated microglial states. CHPV infection resulted in marked downregulation of homeostatic genes, including *Sall1, P2ry12, Tmem119, Cx3cr1, Hexb* and *Mef2a*, accompanied by selective induction of disease-associated/inflammatory genes such as *Ccl2* and *Lgals3*. In contrast, several canonical DAM-associated genes, including *Trem2*, remained decreased (**Fig. 2F**). Thus, CHPV infection induced substantial remodeling of the microglial transcriptional state, characterized by loss of homeostatic identity and acquisition of an inflammatory, but non-canonical DAM-like phenotype.

Pathway enrichment revealed distinct cell-type-specific responses across the selected four populations. CGNs showed enrichment of synaptic signaling, MHC-I antigen presentation and type I IFN pathways, indicating an antiviral neuronal response (**Fig. 2G**). OB-out GABAergic neurons were characterized predominantly by growth, proliferation, developmental and inflammatory wound-healing pathways, with suppression of antiviral response programs (**Fig. 2H**). Microglia were enriched for cell migration, adhesion, proliferation and antiviral defense pathways, consistent with an immune-responsive state (**Fig. 2I**). Astrocytes on the other hand showed reduced anti-viral defense (**Fig. 2J**). Overall, these findings indicate cell-type-specific remodeling of antiviral, neuronal and inflammatory pathways following CHPV infection. In particular, the selective antiviral response of cerebellar granule neurons and the divergent suppression of antiviral programs in astrocytes and selective neuronal populations suggest that CNS cell types differ markedly in their response to CHPV infection.

### CHPV-induced neuronal death is preceded by early disruption of mitochondrial antiviral response and a delayed interferon response

The single-cell transcriptomic analysis revealed cell type-specific alterations in antiviral, neuronal and inflammatory pathways following CHPV infection, prompting us to investigate the early molecular events underlying this response. Since mitochondria serve as central regulators of antiviral signaling during RNA virus infection through the RIG-I-MAVS axis [14, 15], we next examined whether CHPV infection perturbs mitochondrial integrity during early stages of neuronal infection. Assessment of mitochondrial membrane potential using TMRM staining demonstrated a progressive reduction in mitochondrial membrane potential in CHPV-infected SH-SY5Y cells compared to mock-infected controls, as observed by both immunofluorescence imaging and flow cytometric analysis (**Fig. 3A, B**). Given the central role of MAVS in mitochondrial antiviral signaling [40], we next investigated whether CHPV infection alters MAVS protein levels. Immunoblot analysis demonstrated a progressive reduction in MAVS protein levels in both CHPV-infected SH-SY5Y cells and in the brain lysates from CHPV-infected mice (**Fig. 3C-D**), indicating that disruption of mitochondrial antiviral signaling occurs both *in vitro* and *in vivo* during CHPV neuropathogenesis. MAVS serves as the central signaling hub downstream of the cytosolic RNA sensor RIG-I, where it assembles a multiprotein signalosome that recruits adaptor proteins, including TRAF3 and TRAF6, to activate the TBK1-IRF3/IRF7 and NF-κB pathways [41]. This signaling cascade culminates in the production of type I/III IFNs and the induction of IFN-stimulated genes (ISGs), which are essential for mounting an effective antiviral response. To determine whether disruption of MAVS integrity alters the intrinsic antiviral response of infected neurons, we performed bulk RNA sequencing (RNA-Seq) (#GSE346794) of CHPV-infected SH-SY5Y cells at 4, 8 and 24 hpi and compared the data with mock-infected controls. Differential expression analysis (log₂FC ≥ 1, FDR ≤ 0.05) revealed extensive transcriptional reprogramming throughout infection, with 1,723 upregulated and 999 downregulated genes at 4 hpi, 1,697 upregulated and 1,308 downregulated genes at 8 hpi, and a marked increase to 2,178 upregulated and 2,456 downregulated genes at 24 hpi, (**Fig. S2A**). Heatmap analysis showed only minimal induction of key anti-viral signaling molecules, including *TBK1, IRF3, IRF7* and components of the JAK-STAT pathway (*STAT1, STAT2, STAT3, JAK1, JAK2, and JAK3*) at 4 and 8 hpi (**Fig. 3E**). Consistent with this, transcription of type I and type III interferons was markedly attenuated at early time points, with IFNB and IFNL family members (*IFNL1, IFNL2, IFNL3*) exhibiting low or negligible expression at 4 and 8 hpi (**Fig. 3E**). In contrast, a robust induction of both type I and type III interferons was observed at 24 hpi (**Fig. 3E**), which was further validated by qRT-PCR analysis (**Fig. 3F, G**). Additionally, immunoblot analysis of the downstream effectors of IFN signalling showed increased phosphorylation of TBK1, IRF3 and IRF7 at later time points of infection (16 and 24 hpi) (**Fig. 3H**), which coincided with elevated IFN responses, validating the transcriptomic analysis.

**Figure 3.**
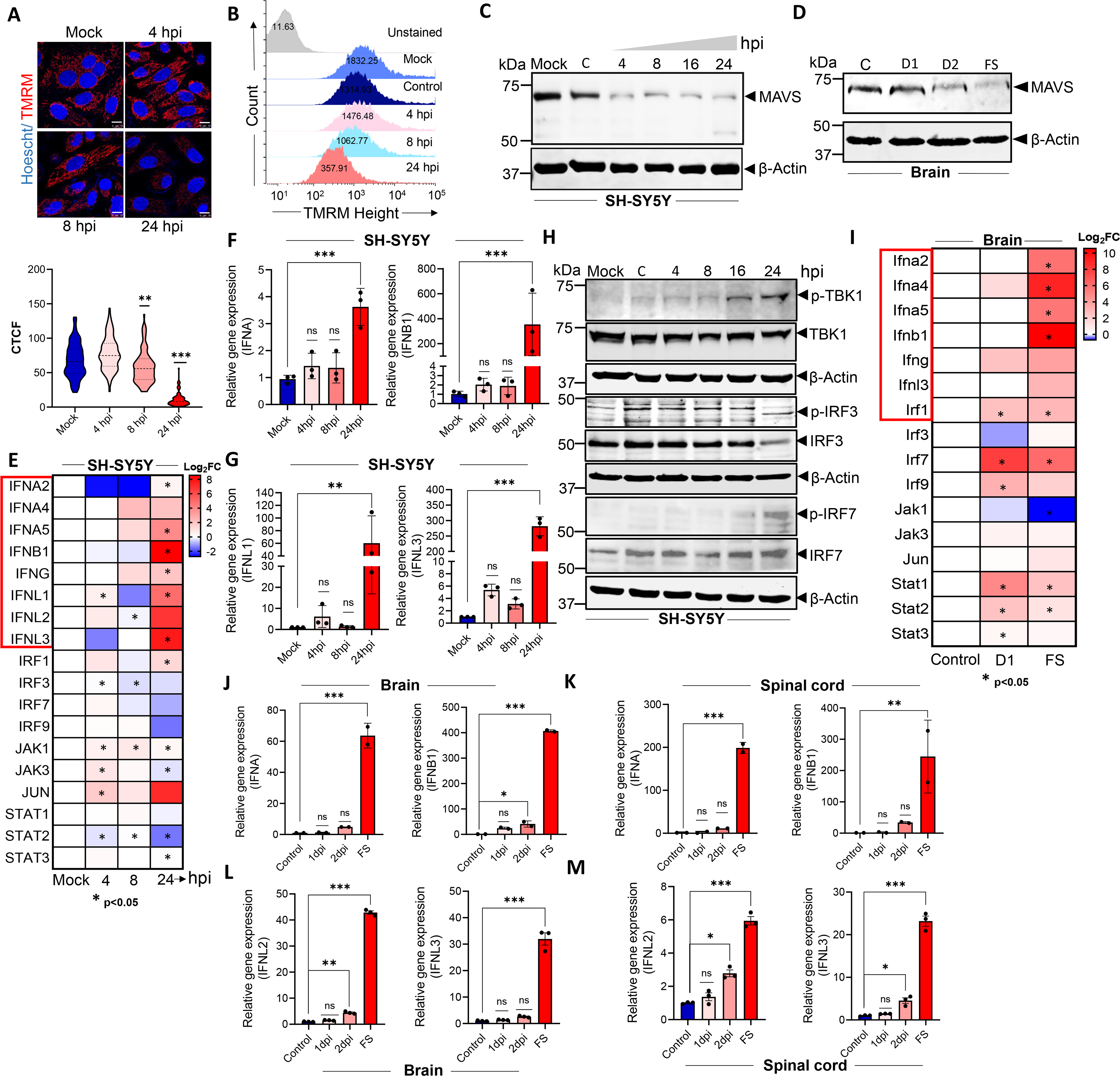
Early disruption of mitochondrial antiviral response precedes CHPV-induced neuronal death that is accompanied by a delayed interferon response. (A) (Top panel) Representative confocal images of SH-SY5Y cells incubated with 100 nM TMRM (red) and Hoechst 33342 (blue) to assess mitochondrial membrane depolarization following CHPV infection (MOI-0.1) and analyzed at 4, 8 and 24 hpi. (Scale = 10 μm). (Bottom panel) Corrected total cell fluorescence (CTCF) values of TMRM intensity of mock vs CHPV-infected cells. **(B)** Flow cytometric analysis of mitochondrial membrane potential using TMRM. The representative histogram analysis shows TMRM fluorescence in SH-SY5Y cells following CHPV infection (MOI-0.1) and analyzed at 4, 8 and 24 hpi. **(C)** Immunoblot analysis of MAVS protein levels in the whole-cell extracts of SH-SY5Y cells at 4, 8, 16 and 24 hpi and compared with mock-infected and control (C; uninfected) cells. β-Actin was used as a loading control. **(D)** Immunoblot analysis of MAVS protein levels in mouse brain lysates at D1, D2 and FS stages post-CHPV infection and compared with control. β-Actin was used as a loading control. **(E)** Heatmap showing relative mRNA expression of differentially expressed genes of the interferon pathway obtained from bulk RNA sequencing in CHPV-infected SH-SY5Y cells at 4, 8 and 24 hpi and compared with mock-infected cells (#GSE346794). *Represent the significant genes (p≤0.05). **(F)** qRT-PCR analysis of the relative gene expression of type-I interferons (IFNA and IFNB1) in SH-SY5Y cells post-CHPV infection. **(G)** qRT-PCR analysis of the relative gene expression of type-3 interferons (IFNL1 and IFNL3) in SH-SY5Y cells post-CHPV infection. **(H)** Immunoblot analysis of pTBK1/TBK1, pIRF3/IRF3, pIRF7/IRF7 protein levels in the whole-cell extracts of CHPV-infected SH-SY5Y cells at 4, 8 and 24 hpi and compared with mock-infected and control cells. β-Actin was used as a loading control. **(I)** Heatmap showing relative mRNA expression of differentially expressed genes of the interferon pathway obtained from bulk RNA sequencing in mouse brain tissue (GSE). *Represent the significant genes (p≤0.05). **(J)** qRT-PCR analysis of the relative gene expression of type-I interferons (IFNA and IFNB1) in mouse brain tissue post-CHPV infection. **(K)** qRT-PCR analysis of the relative gene expression of type-I interferons (IFNA and IFNB1) in mouse spinal cord tissue post-CHPV infection. **(L)** qRT-PCR analysis of the relative gene expression of type-III interferons (IFNL2 and IFNL3) in mouse brain tissue post-CHPV infection. **(M)** qRT-PCR analysis of the relative gene expression of type-III interferons (IFNL2 and IFNL3) in spinal cord tissue post-CHPV infection. Data are presented as mean ± SD. Statistical significance was determined by one-way ANOVA with post hoc Tukey’s multiple comparison test (**P < 0.01; ***P < 0.001; n=3 biological replicates).

Bulk RNA sequencing of mouse brains (#GSE347261) showed that at D1, 6,923 genes were significantly upregulated, whereas only 504 genes were downregulated, indicating a predominant activation of host gene expression early during infection. By FS, the transcriptional response became more extensive, with 9,689 upregulated and 4,062 downregulated genes, reflecting widespread reprogramming of host transcriptional pathways during disease progression (**Fig. S2B**). Genes associated with canonical IFN signaling pathways also exhibited limited induction at early stages of infection (D1) in CHPV-infected BALB/c mice (**Fig. 3I**). Consistent with this, qRT-PCR validation in brain tissue confirmed that type I and III IFN genes remained weakly expressed during early infection, with a significant upregulation observed only during the later period of infection when the symptoms appeared (**Fig. 3J, L**). A similar temporal pattern was observed in the spinal cord, where qRT-PCR analysis showed delayed induction of IFN genes despite increasing viral load (**Fig. 3K, M**). Consistent with this, transcriptomic analysis of CHPV-infected neuronal cells and mouse brain revealed a robust induction of ISGs predominantly during the late stage of infection (**Fig. S2C-D**). Together, these findings indicate a delayed activation of the interferon-mediated antiviral response during CHPV infection.

### Pharmacological inhibition of lysosomal function prevents reduction of MAVS protein levels following CHPV infection

There is accumulating evidence that MAVS is degraded through multiple mechanisms [42]. Importantly, MAVS is targeted for autophagy-lysosome dependent degradation as a key mechanism of virus-mediated immune evasion [43].To determine whether the lysosomal pathway contributes to MAVS degradation following CHPV infection, SH-SY5Y cells were pretreated for 1 h with either chloroquine (CQ), an inhibitor of lysosomal acidification, or bafilomycin A1 (Baf), which blocks autophagosome-lysosome fusion [44], prior to CHPV infection (**Fig. S3A**). Compared to CHPV-infected SH-SY5Y cells, both treatments markedly attenuated MAVS degradation, sustaining its steady-state protein levels at 24 hpi (**Fig. S3A**). Consistent with these findings, LAMP2 staining revealed increased colocalization of MAVS with lysosomes during CHPV infection starting at 4 hpi (**Fig. S3B**), further supporting the involvement of the lysosomal pathway in MAVS turnover. Notably, inhibition of lysosomal function by CQ also attenuated CHPV-induced cell death along with reduction in viral replication and release (**Fig. S3C-E**), indicating that lysosome-dependent processes are critical mediators of CHPV-induced reduction in MAVS protein levels and immune evasion strategy. Given prior reports that CQ can inhibit viral entry [45], we next sought to determine whether the protective effects of CQ in our system were attributable to impaired CHPV entry. To address this, virions were pre-labeled with the Paul Karl Horan (PKH) dye, washed extensively to remove excess dye and subsequently used to infect SH-SY5Y cells (**Fig. S3F**). PKH fluorescence was readily detected within infected cells even in the presence of CQ at 4 hpi, indicating that viral entry was not blocked under these conditions (**Fig. S3F**). In addition, CQ pre-treatment markedly attenuated the induction of type-I IFN response (IFNA and IFNB1) in CHPV-infected cells (**Fig. S3G-H**). Collectively, these data demonstrate that lysosomal activity contributes to the loss of MAVS protein abundance during CHPV infection and thereby negatively control host antiviral response. Further, the persistence of TBK1 and IRF3/7 phosphorylation despite MAVS loss revealed an unexpected dissociation between mitochondrial antiviral platform integrity and downstream interferon signaling, pointing towards the engagement of alternative or MAVS-independent anti-viral signaling mechanisms during CHPV infection.

### CHPV infection is associated with heightened inflammatory response and pyroptotic cell death

So far, our findings indicate that despite substantial MAVS depletion, CHPV-infected neurons retain downstream antiviral signaling, as evidenced by persistent TBK1 and IRF3/7 phosphorylation. This suggests that CHPV infection may engage an alternative, MAVS-independent pathway to sustain downstream IFN signaling. Since activation of these pathways is often accompanied by the production of inflammatory mediators, we next wanted to examine the temporal dynamics of the inflammatory response associated with CHPV infection and subsequent neuronal cell death. Acute viral infections, particularly in encephalitic models, are known to trigger robust inflammatory responses as part of the host antiviral defense [46, 47]. While these responses are essential for viral clearance, their dysregulation or persistence can contribute to severe disease pathology. To determine the host inflammatory responses during CHPV infection, we assessed the expression of pro-inflammatory mediators in CHPV-infected SH-SY5Y cells and mouse brain tissue using bulk RNA-Seq analysis, followed by qRT-PCR validation. Interestingly, the transcript levels of the pro-inflammatory cytokine *IL1B* increased at 4 hpi, followed by a decline as the infection progressed in SH-SY5Y cells (**Fig. 4A, C**). In contrast, *TNFA* and *IL6* expressions displayed a dynamic temporal pattern, with reduced expression at 8 hpi followed by a marked transcriptional activation at 24 hpi (**Fig. 4A, C**). Next, we quantified the protein levels of these cytokines in the culture supernatants of CHPV-infected SH-SY5Y cells by ELISA. As similar to its transcript levels, CHPV infection induced a transient increase in IL1β secretion, which peaked at 4 hpi and progressively declined to near basal levels by 24 hpi (**Fig. 4D**). In contrast, TNFα secretion was initially suppressed following infection, whereas IL6 secretion showed modest induction during the early phase of infection but a pronounced increase at 24 hpi (**Fig. 4D**). Consistent with these observations, bulk RNA-Seq and qRT-PCR analyses of CHPV-infected mouse brain and spinal cord tissue revealed a sustained inflammatory response beginning at D1 and markedly increasing at FS stage, coinciding with peak disease severity (**Fig. 4B, E and F**). Notably, unlike neuronal cells, CHPV-infected mouse brain tissue exhibited sustained *Il1B* expression throughout the course of infection, suggesting the contribution of additional cellular sources within the brain. To explore this possibility, we next examined the inflammatory response in microglial N9 cells following CHPV infection at an MOI of 0.1. However, there was no induction of the proinflammatory cytokines following CHPV infection (**Fig. 4G**), possibly due to a lack of productive viral replication within these cells as compared to the neuronal SH-SY5Y cells. However, since our data indicated a heightened inflammatory response within the brain microglial cells, we infected the N9 cells at a higher MOI of 1 and observed increased inflammatory responses in these cells, especially *IL1B*, which indicates that microglia could contribute to the higher *Il1B* in the bulk RNA-Seq in the CHPV-infected brains as compared to SH-SY5Y cells (**Fig. 4H**). Together, our data suggest that while neurons constitute the primary site of CHPV replication and initiate early inflammatory signaling, microglia are likely recruited into the inflammatory cascade as viral burden increases, thereby amplifying the neuroinflammatory environment during the later stages of infection. This observation is consistent with recent evidence demonstrating that the outcome of CHPV infection is profoundly influenced by infection multiplicity, with high MOI conditions favoring extensive cell death during productive infection [48].

**Figure 4.**
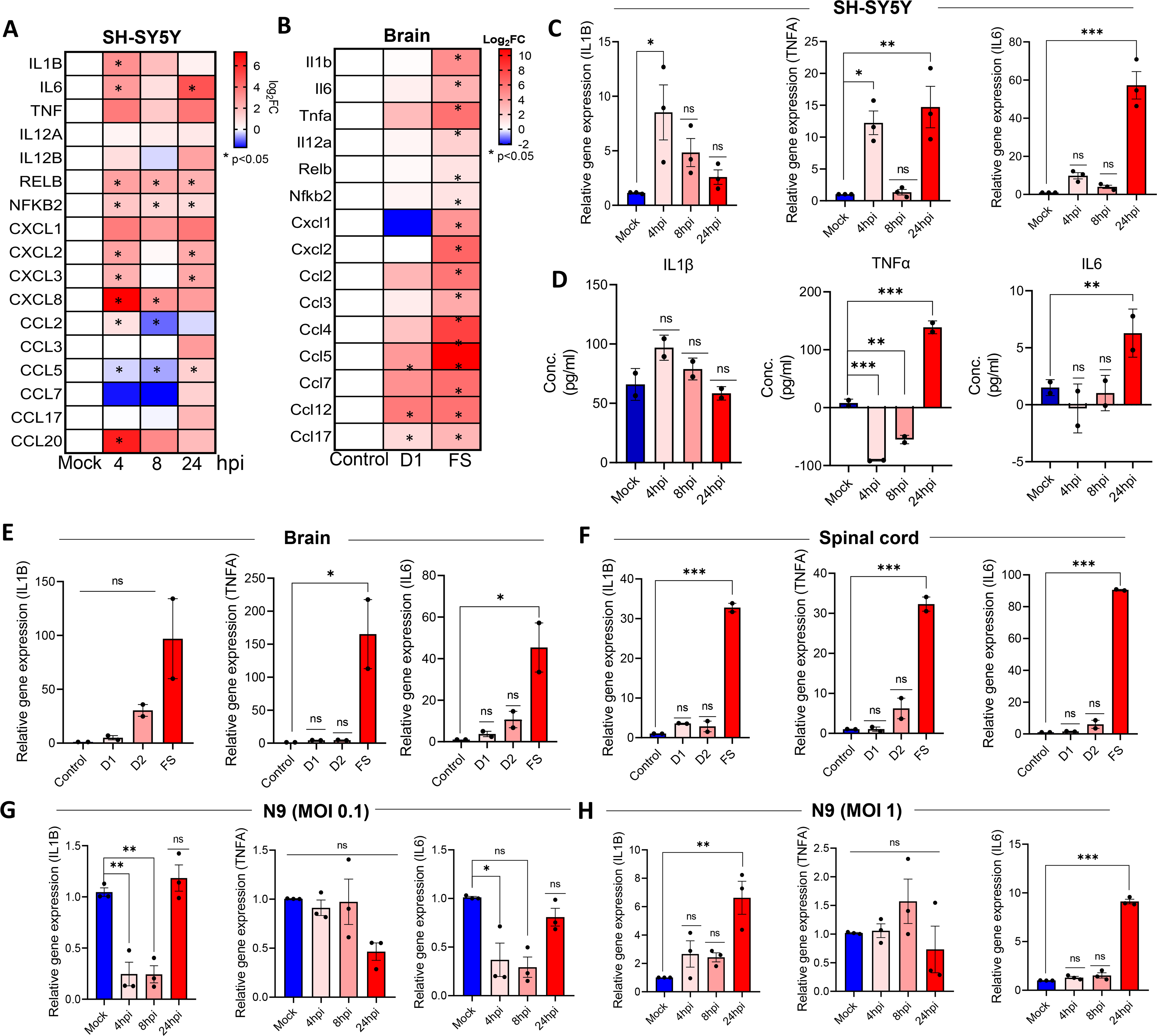
CHPV-induced cell death is accompanied by heightened inflammatory response during the late stage of infection. **(A)** Heatmap showing relative mRNA expression of differentially expressed genes of the inflammatory response pathway obtained from bulk RNA sequencing in CHPV-infected SH-SY5Y cells at 4, 8 and 24 hpi and compared with mock-infected cells (GSE). * Represents the significant genes (p≤0.05). **(B)** Heatmap showing relative mRNA expression of differentially expressed genes of the inflammatory response pathway obtained from bulk RNA sequencing in mouse brain tissue (GSE). *Represent the significant genes (p≤0.05). **(C)** qRT-PCR analysis of the relative gene expression of inflammatory genes (IL1B, TNFA and IL6) in SH-SY5Y cells post-CHPV infection. **(D)** ELISA assay of pro-inflammatory cytokines IL1β, TNFα and IL6 in cell supernatant collected from SH-SY5Y cells at different time points post CHPV infection. (n=2). **(E)** qRT-PCR analysis of the relative gene expression of inflammatory genes (IL1B, TNFA and IL6) in mouse brain tissue post-CHPV infection. **(F-H)** qRT-PCR analysis of the relative gene expression of inflammatory genes (IL1B, TNFA and IL6) in mouse spinal cord tissue post CHPV infection (F) and in N9 cells post-CHPV infection (MOI-0.1) (G) and (MOI-1) (H). Data are presented as mean ± SD. Statistical significance was determined by one-way ANOVA with post hoc Tukey’s multiple comparison test (**P < 0.01; ***P < 0.001; n=3 biological replicates).

Since CHPV induced robust cell death in neuronal SH-SY5Y cells, we sought to delineate the underlying mode of cell death. Among the major forms of programmed cell death, that includes apoptosis, necrosis, autophagy and pyroptosis, the latter has emerged as a critical mediator of inflammation-associated tissue injury [49]. Pyroptosis is a lytic form of programmed cell death characterized by the formation of plasma membrane pores by members of the GSDM family following proteolytic cleavage by activated caspases [50, 51]. In the canonical pathway, inflammasome activation leads to caspase-1-mediated cleavage of GSDMD, whereas activation of apoptotic caspase-3 can cleave GSDME, converting apoptotic signaling into secondary pyroptosis [52, 53]. Both pathways culminate in membrane rupture and the release of pro-inflammatory mediators, including IL1β and IL18, thereby amplifying local inflammation and tissue damage [54]. While CHPV has been extensively reported to induce neuronal apoptosis, whether inflammatory pyroptosis, contributes to CHPV-induced neuronal injury has not been reported yet. Transcriptomic analysis of genes associated with programmed cell death revealed selective induction of inflammatory cell death pathways during CHPV infection. In SH-SY5Y cells, modest transcriptional upregulation of pyroptosis-associated genes was observed, whereas genes associated with the intrinsic apoptotic pathway exhibited relatively limited changes (**Fig. 5A**). Similar but more pronounced upregulation of pyroptosis-associated genes, was also detected in infected mouse brain tissue (**Fig. 5B**), suggesting amplification of inflammatory cell death pathways within the infected CNS. Since most components of the apoptotic and pyroptotic pathways are primarily regulated through post-translational proteolytic processing rather than transcriptional induction, we next examined the activation status of key apoptotic and pyroptotic execution proteins by immunoblotting. CHPV infection induced a moderate increase in NLRP3 protein levels, accompanied by cleavage of the inflammatory caspase-1 with maximal activation observed at 24 hpi (**Fig. S4A**). Surprisingly, despite robust caspase-1 activation, cleavage of GSDMD into its canonical pore-forming N-terminal fragment (∼31–35 kDa) was not detected throughout the course of infection. Instead, we consistently observed accumulation of a lower molecular weight GSDMD fragment of approximately 25 kDa specifically at 24 hpi (**Fig. S4B**). A similar ∼25 kDa GSDMD fragment has previously been reported during *Leishmania* infection [55], where it was proposed to represent an inhibitory cleavage product incapable of mediating pyroptotic pore formation. These findings indicate that although CHPV activates the inflammasome, canonical GSDMD-dependent pyroptosis is probably not efficiently executed in CHPV-infected SH-SY5Y cells and needs further investigation. We next investigated whether CHPV infection engages the caspase-3/GSDME axis, an alternative pathway capable of mediating secondary pyroptosis [32]. We observed that CHPV infection induced progressive cleavage of caspase-8, caspase-3 and its substrate PARP1 (**Fig. 5C**) as well as caspase-9 (**Fig. S4C**), accompanied by proteolytic processing of GSDME and accumulation of its pore-forming N-terminal fragment, which became most prominent at 24 hpi (**Fig. 5C**). CHPV infection of SH-SY5Y cells also resulted in a significant increase in extracellular lactate dehydrogenase (LDH) release (**Fig. 5D**), indicating CHPV infection leads to a progressive loss of plasma membrane integrity, a hallmark of pyroptotic cell death. Consistent with this finding, morphological examination of CHPV-infected SH-SY5Y cells by scanning electron microscopy (SEM) revealed marked cellular swelling, membrane ballooning and extensive disruption of plasma membrane integrity with visible pores in the membrane by 24 hpi compared with mock-infected cells, features characteristic of pyroptotic cell death (**Fig. 5E**). Consistent with the observations in cultured SH-SY5Y cells, infected brains exhibited robust cleavage of GSDME at FS (**Fig. 5F**), indicating that GSDME-mediated inflammatory cell death may accompany CHPV-induced neuropathogenesis *in vivo* as well.

**Figure 5.**
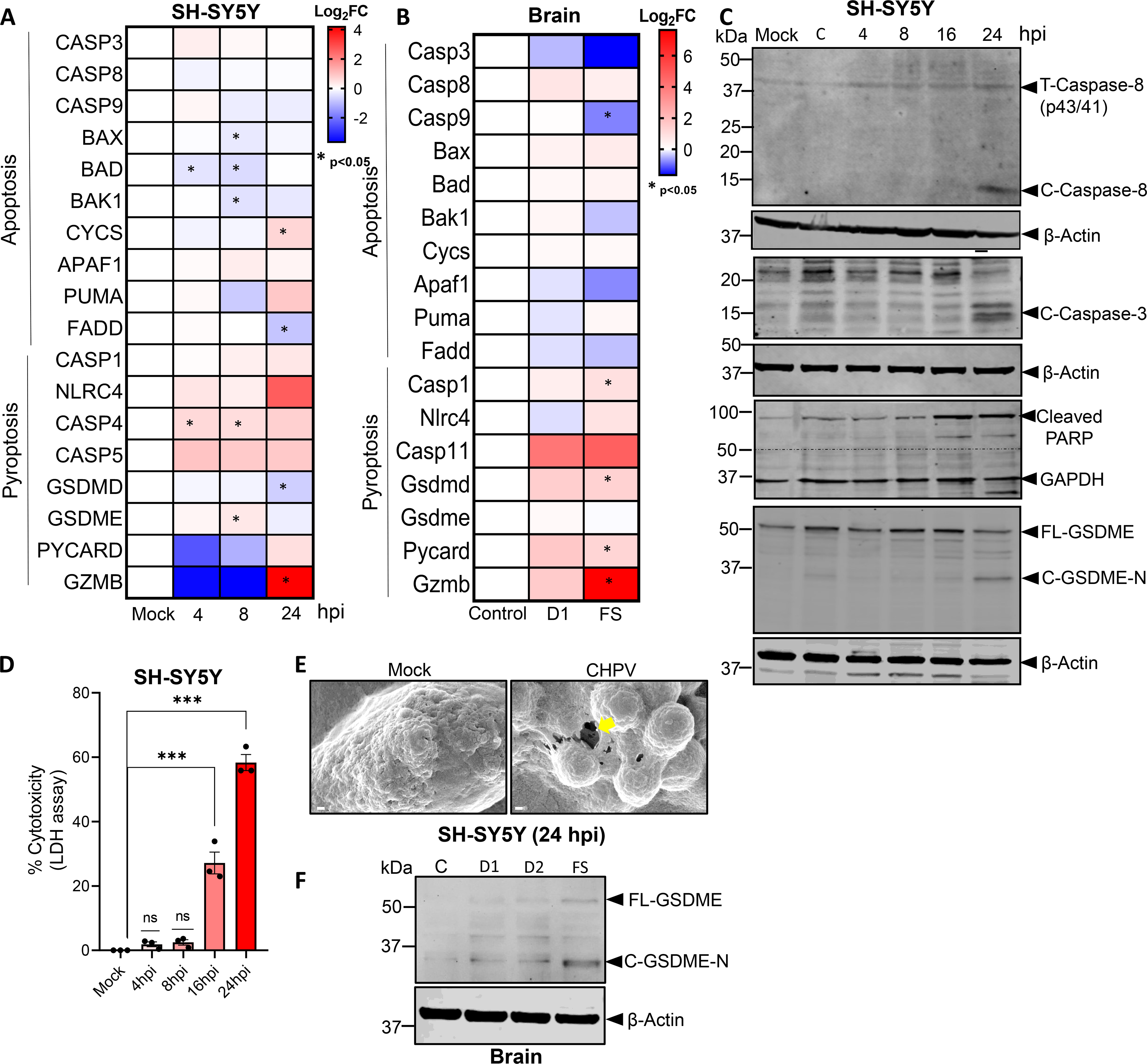
CHPV-induced inflammatory responses promote pyroptotic cell death in neuronal SH-SY5Y cells. **(A,B)** Heatmap showing relative mRNA expression levels of differentially expressed genes of cell death pathways at indicated time points obtained from bulk RNA sequencing in (A) SH-SY5Y cells and (B) in mouse brain tissue. * Represents the significant genes (p≤0.05). **(C)** Immunoblot analysis of caspase 8 (cleaved), cleaved caspase 3, cleaved PARP, total and cleaved GSDME-N terminal (GSDME) protein levels in the whole-cell extracts of CHPV-infected SH-SY5Y cells at 4, 8, 16 and 24 hpi and compared with mock-infected and control (C) cells. β-Actin and GAPDH was used as a loading control. **(D)** Cell cytotoxicity measurement by lactate dehydrogenase (LDH) assay in SH-SY5Y cells at indicated time points post-CHPV infection. **(E)** Scanning electron microscope (SEM) images representing SH-SY5Y cell membrane post-CHPV infection. **(F)** Immunoblot analysis of total and cleaved GSDME (N terminal) protein levels in mouse brain lysates at D1, D2 and FS. β-Actin was used as a loading control. The yellow arrows represent the pores at 24 hpi. (Scale bar = 200 nm). Data are presented as mean ± SD. Statistical significance was determined by one-way ANOVA with post hoc Tukey’s multiple comparison test (**P < 0.01; ***P < 0.001; n=3 biological replicates).

Together, these findings indicate that CHPV infection in neuronal SH-SY5Y cells induces a GSDME-mediated inflammatory cell death program, where unlike classical pyroptotic program, GSDME acts as a bridge connecting the apoptotic machinery to lytic cell death.

### STING activation drives inflammation-associated pyroptotic cell death during CHPV infection

Since MAVS was degraded during early infection, we sought to determine the host factors responsible for triggering inflammation and subsequent pyroptotic cell death program. The cGAS-STING pathway is known to induce an antiviral state by stimulating the production of both IFNs and pro-inflammatory cytokines [56]. cGAS is known to bind cytosolic dsDNA, leading to the production of cGAMP and subsequent activation of STING signaling. Upon activation, STING translocate from the endoplasmic reticulum to the Golgi apparatus, where it recruits and activates TBK1, resulting in phosphorylation of IRF3/7 and induction of type I IFN and pro-inflammatory gene expression [57]. However, in contrast to DNA viruses, the engagement of the cGAS-STING pathway in the context of RNA virus infection is not fully understood. To determine whether CHPV infection can activate cGAS-STING pathway and downstream signaling, we first checked the STING-phosphorylation pattern in both *in vitro* and *in vivo* models. STING was phosphorylated in both CHPV-infected SH-SY5Y cells as well as mouse brain tissue, wherein the timing of phosphorylation coincided with the induction of type-I/III IFNs and pro-inflammatory cytokines (**Fig. 6A-B**). To further assess STING activation, we examined its subcellular localization in SH-SY5Y cell lines using immunofluorescence analysis (**Fig. 6C**). In mock-infected cells, STING predominantly co-localized with the ER marker calreticulin, displaying an ER distribution pattern consistent with its basal state (**Fig. 6C Upper and Lower panel**). In contrast, CHPV infection induced a redistribution of STING into discrete punctate structures with reduced overlap with calreticulin. Notably, STING-positive puncta exhibited strong co-localization with the cis-Golgi marker GM130, indicating ER-to-Golgi translocation of STING in response to CHPV infection (**Fig. 6C Middle and Lower panel**).

**Figure 6.**
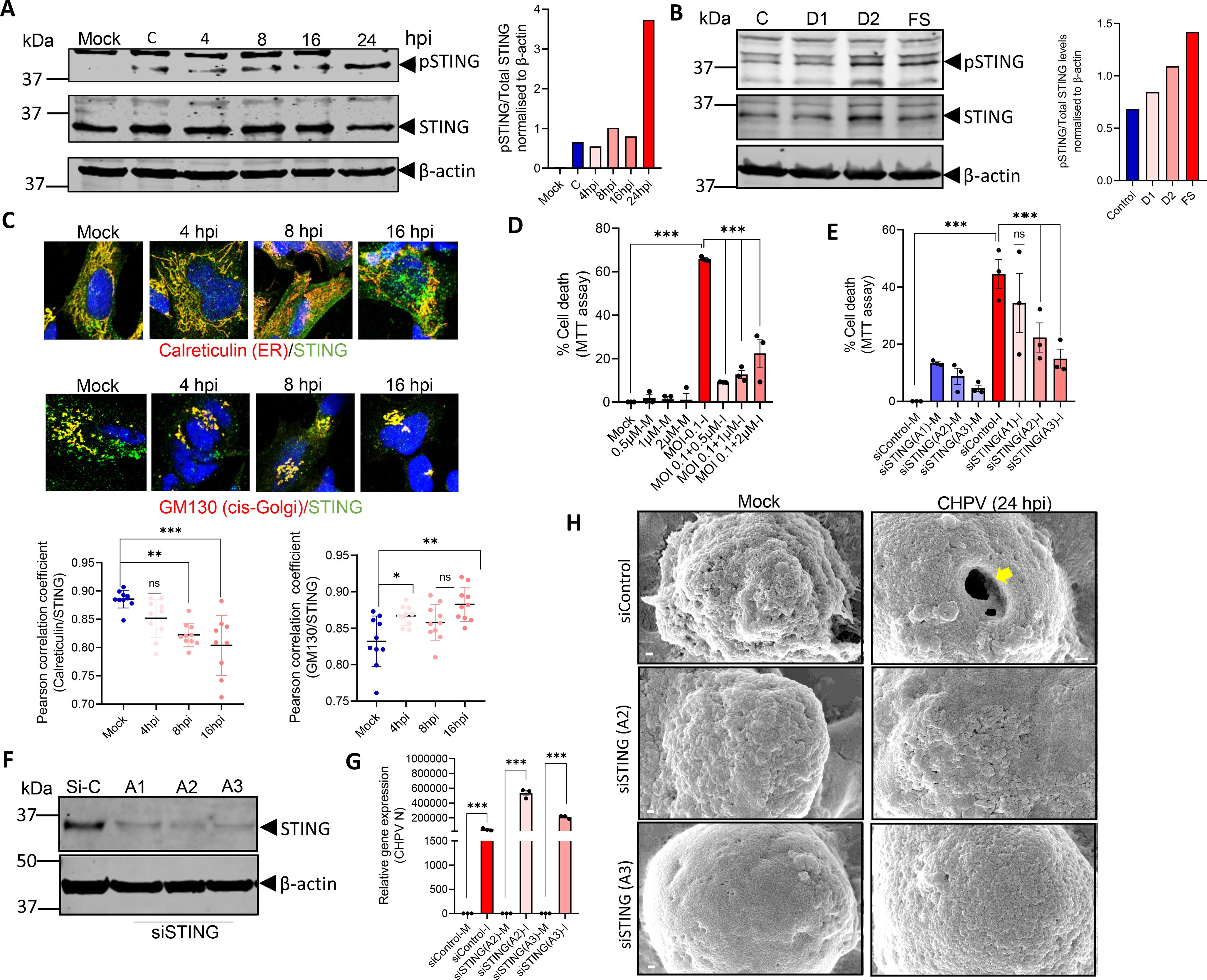
STING activation drives inflammation-associated pyroptotic cell death during CHPV infection. **(A)** (Left panel) Immunoblot analysis of pSTING/STING protein levels in the whole-cell extracts of SH-SY5Y cells at 4, 8, 16 and 24 hpi and compared with mock-infected and control (uninfected) cells. β-Actin was used as a loading control. (Right panel) Quantification of pSTING levels against total STING and β-Actin levels. **(B)** (Left panel) Immunoblot analysis of pSTING/STING protein levels in mouse brain lysates at D1, D2 and FS stages post-CHPV infection and compared with control. β-Actin was used as a loading control. (Right panel) Quantification of pSTING levels against total STING and β-Actin levels. **(C)** Representative confocal images of SH-SY5Y cells post-CHPV infection stained against (Upper panel) Calreticulin (red), STING (green) and DAPI (blue) and (Middle panel) GM130 (red), STING (green) and DAPI (blue) alongwith their PCC value (Lower panel). (Scale bar = 10 µm). **(D)** Percentage of cell death measured by MTT assay in SH-SY5Y cells post-CHPV infection (MOI-0.1) at 24 hpi pretreated with the STING inhibitor STIN-IN-2. **(E)** Percentage of cell death measured by MTT assay in siSTING-transfected SH-SY5Y cells post-CHPV infection (MOI-0.1) at 24 hpi or in cells pretreated with STIN-IN-2 (STING inhibitor) for 1 h. (M=Mock and I=Infected). **(F)** Immunoblot analysis of STING protein levels in the whole-cell extracts of SH-SY5Y cells transfected for 48 h with siSTING. β-Actin was used as a loading control. **(G)** qRT-PCR analysis of the relative gene expression of CHPV N gene in SH-SY5Y cells transfected with siSTING for 48 h and analyzed at 24 hpi. (M=Mock and I=Infected). **(H)** SEM images representing SH-SY5Y cell membrane rupture in siControl vs siSTING-transfected SH-SY5Y cells post-CHPV infection. The yellow arrows represent the pores at 24 hpi. (Scale bar = 200 nm). Data are presented as mean ± SD. Statistical significance was determined by one-way ANOVA with post hoc Tukey’s multiple comparison test (**P < 0.01; ***P < 0.001; n=3 biological replicates).

Since STING relocalization to Golgi attributes to its activation and supports engagement of downstream signaling pathways, we sought to determine the significance of STING activation on CHPV-induced neuronal death. To this end, we pre-incubated SH-SY5Y cells with increasing dose of the STING inhibitor, STIN-IN-2 for 24 h prior to infection (**Fig. 6D**). Treatment with the STING inhibitor markedly reduced CHPV-induced cell death (**Fig. 6D**). To exclude potential pleiotropic effects of pharmacological inhibition and further validate the role of STING in CHPV-induced neuronal death, we used three independent STING-specific siRNAs to knockdown STING in SH-SY5Y cells and subsequently assessed CHPV-induced cellular responses (**Fig. 6E-H**). In contrast to siControl-transfected SH-SY5Y cells, STING knockdown with siSTING-A2 and siSTING-A3 markedly reduced CHPV-induced cell death (**Fig. 6E**), further establishing a critical role for STING signaling in CHPV-associated cytotoxicity. Among the three STING-specific siRNAs, siSTING-A1 produced a modest reduction in CHPV-induced cell death, although this effect did not reach statistical significance; notably, all three siRNAs effectively reduced STING protein levels. Henceforth, we used the siSTING-A2 and siSTING-A3 for further functional analysis (**Fig. 6G-H**). Notably, STING knockdown did not reduce overall viral burden within the cells but prevented the induction of both type-I IFN and inflammatory responses in CHPV-infected SH-SY5Y cells (**Figs. 6G and S5A-E**). In addition, our SEM imaging results also demonstrated that STING KD prevented CHPV-induced morphological alterations and pyroptosis-associated cellular features and pore formation (**Fig. 6H**). Together, these results identify STING as a central regulator of the host inflammatory response to CHPV infection, functioning independently of viral replication and suggesting that STING-mediated immune signaling, rather than viral burden per se, contributes to downstream neuronal pathology.

### CHPV-induced mitochondrial dysfunction promotes mtROS generation, mtDNA release and GSDME association with mitochondria

The cGAS–STING pathway is classically activated by cGAS-mediated recognition of cytosolic double-stranded DNA (dsDNA), a well-established mechanism in response to DNA virus infections [58]. However, the source of cytosolic dsDNA during RNA virus infections has not been studied in-depth, particularly for encephalitic viruses [59]. Given our observation of mitochondrial dysfunction and MAVS depletion accompanied with heightened inflammation during CHPV infection, we hypothesized that mitochondrial integrity may be compromised, leading to the release of mitochondrial DNA (mtDNA) into the cytosol. Such cytosolic mtDNA could serve as a ligand for cGAS, thereby driving STING activation. Supporting this possibility, previous studies have shown that mitochondrial damage, particularly under conditions of oxidative stress and elevated reactive oxygen species (ROS), can promote the mtDNA release into the cytoplasm [60, 61]. To test this hypothesis, we first checked the levels of mtROS in CHPV-infected SH-SY5Y cells (**Fig. 7A-B**). FACS analysis using the live-cell permeable mitochondrial superoxide indicator MitoSOX Red revealed a progressive increase in mtROS levels following CHPV infection, reaching a maximum at 24 hpi (**Fig. 7A**). Consistent with these findings, immunofluorescence analysis independently confirmed the accumulation of mtROS in SH-SY5Y cells at 24 hpi (**Fig. 7B**).

**Figure 7.**
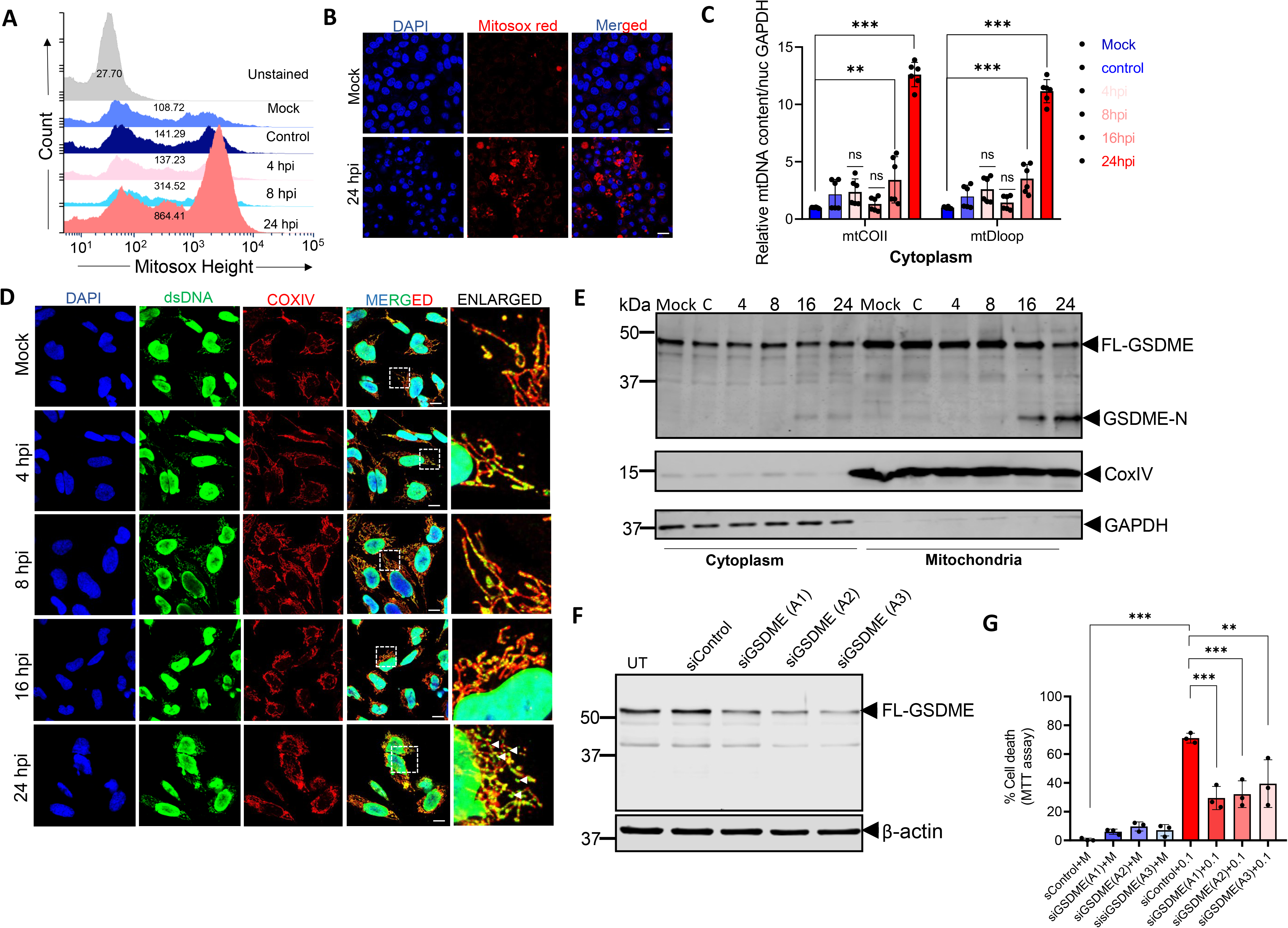
CHPV-induced mitochondrial dysfunction promotes mtROS generation, mtDNA release and GSDME association with mitochondria. **(A)** Flow cytometric analysis of mtROS production using MitoSOX staining. The representative histogram analysis shows MitoSOX fluorescence in mock vs CHPV-infected cells analyzed at the indicated time points. **(B)** Representative confocal images of SH-SY5Y cells stained with MitoSOX dye (red) and Hoeschst 33342 (blue) in mock vs CHPV-infected cells at the indicated time points. (Scale bar = 10 µm). **(C)** qRT-PCR analysis of the relative gene expression of mitochondrial DNA (mtDloop and mtCO2) in the cytoplasmic fraction of SH-SY5Y cells post-CHPV infection. Nuclear GAPDH was used as an internal control. **(D)** Immunofluorescence images of SH-SY5Y cells post-CHPV infection stained against dsDNA (green), COXIV (red) and DAPI (blue). The white arrowheads represent the presence of dsDNA in cellular cytoplasm at 24hpi. (Scale bar = 10 µm). **(E)** Immunoblot analysis of total and cleaved GSDME-N protein levels in cytoplasmic and mitochondrial fraction of SH-SY5Y cells at 4, 8, 16 and 24 hpi. COXIV was used as a loading control for mitochondrial fraction and GAPDH for cytoplasmic fraction. **(F)** Immunoblot analysis of GSDME protein levels in the whole-cell extracts of SH-SY5Y cells transfected for 48 h with siGSDME. β-Actin was used as a loading control. **(G)** Percentage of cell death measured by MTT assay in SH-SY5Y cells post-CHPV infection (MOI-0.1) at 24 hpi in cells transfected with GSDME-specific siRNA for 48 h. (M = mock; I = infected). Data are presented as mean ± SD. Statistical significance was determined by one-way ANOVA with post hoc Tukey’s multiple comparison test (**P < 0.01; ***P < 0.001; n=3 biological replicates).

Given that mitochondrial oxidative stress can promote mitochondrial membrane perturbation and subsequent mtDNA release, we next examined cytosolic mtDNA following CHPV infection. Cytosolic mtDNA progressively increased during infection, reaching a maximum at 24 hpi (**Fig. 7C**). This was further corroborated by immunofluorescence staining with an anti-dsDNA antibody, which revealed enhanced cytosolic accumulation of dsDNA in infected cells at 24 hpi (**Fig. 7D**). Further, CHPV infection resulted in increased TFAM accumulation in the cytosolic fraction (**Fig. S6**), further supporting mitochondrial perturbation and the release of mtDNA into the cytosol during CHPV infection. Although the presence of cytosolic mtDNA and TFAM indicated compromised mitochondrial integrity, it remained unclear whether mtDNA release resulted from regulated mitochondrial membrane permeabilization or nonspecific mitochondrial membrane disruption during CHPV infection. Recent studies have implicated members of the GSDM family in mitochondrial membrane permeabilization under inflammatory conditions [62]. Notably, GSDM proteins beyond GSDMD have been reported to target mitochondria [63], prompting us to examine the subcellular localization of GSDME during CHPV infection (**Fig. 7F**). Subcellular fractionation revealed a progressive accumulation of cleaved GSDME in the mitochondrial fraction, with maximal levels observed at 24 hpi (**Fig. 7F**). To determine whether GSDME contributes functionally to CHPV-induced cytotoxicity, we used three independent GSDME-specific siRNAs to deplete GSDME (**Fig. 7G**). GSDME knockdown markedly reduced CHPV-induced cell death (**Fig. 7H**), supporting a functional role for GSDME in CHPV-induced neuronal injury.

### GSDME promotes mtDNA release and STING activation during CHPV infection

We next asked whether GSDME contributes to mitochondrial damage and subsequent mtDNA release during CHPV infection. Depletion of GSDME using GSDME-specific siRNA, as described above, markedly reduced the accumulation of mtDNA in the cytosolic fraction of CHPV-infected SH-SY5Y cells (**Fig. 8A**). These findings provide causal evidence for a role of GSDME in CHPV-induced mtDNA release, a previously unrecognized link between GSDME activation and mitochondrial DNA mobilization during viral infection. Given the marked reduction in CHPV-induced cell death following GSDME depletion, we next examined whether GSDME knockdown also affected viral replication and downstream antiviral and inflammatory responses. GSDME depletion resulted in a significant reduction in viral load (**Fig. S7A**), accompanied by attenuation of type I IFN (**Fig. S7B-C**) and inflammatory responses (**Fig. S7D-F**). Since GSDME depletion reduced, but did not completely abrogate cytosolic mtDNA accumulation, we next examined whether this was associated with changes in STING activation. Notably, CHPV-infected GSDME knockdown SH-SY5Y cells showed reduced levels of phosphorylated STING at 24 hpi (**Fig. 8B**), linking GSDME-dependent mtDNA release to STING activation during CHPV infection. To further explore the relationship between GSDME and STING, we next examined GSDME activation following STING depletion (**Fig. 8C**). Interestingly, STING knockdown reduced both total GSDME abundance and the levels of its cleaved product following CHPV infection (**Fig. 8C**), suggesting a reciprocal relationship between STING signaling and GSDME activation. The molecular basis and functional significance of this reciprocal regulation remain to be further investigated.

**Figure 8.**
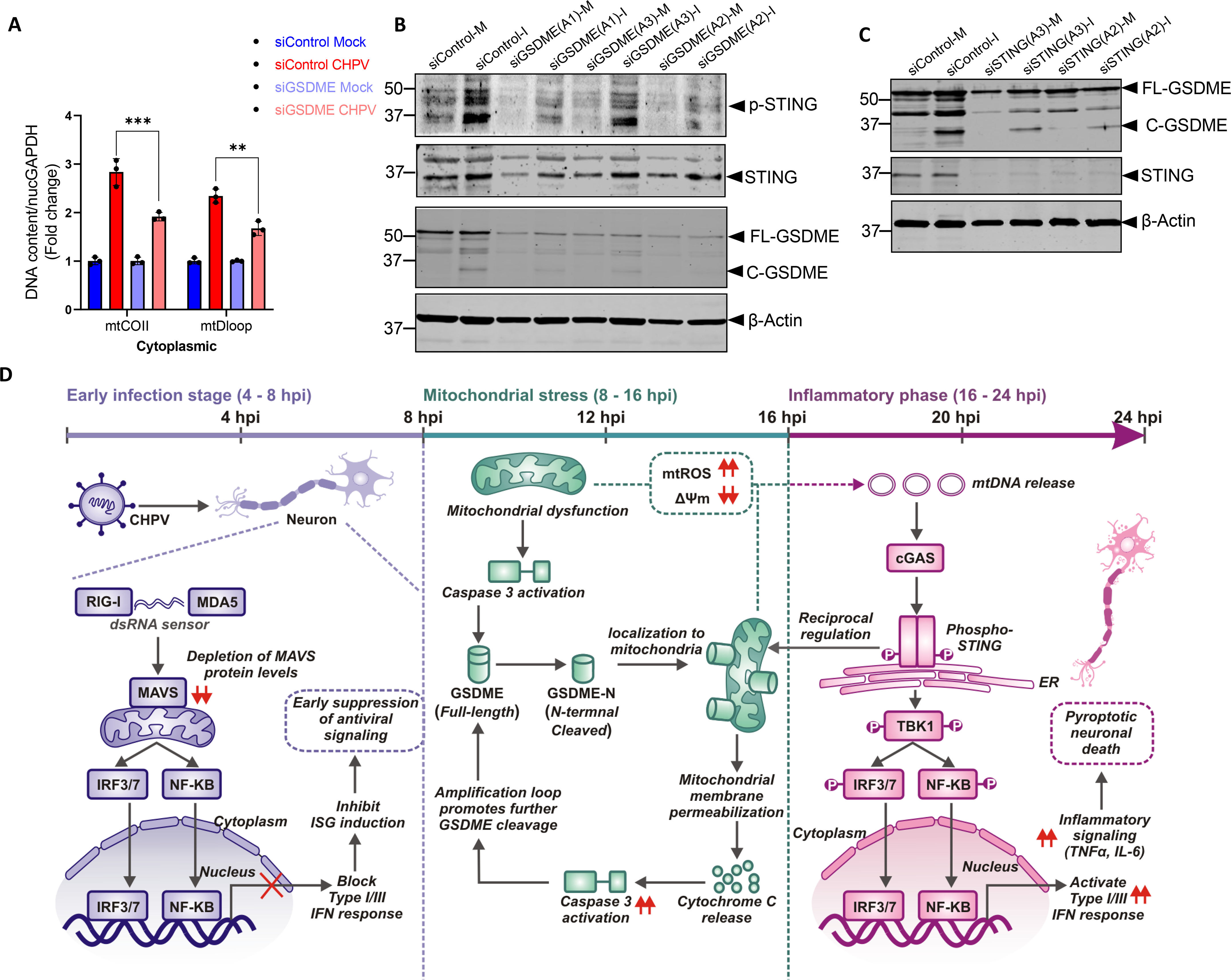
GSDME promotes mtDNA release and STING activation during CHPV infection. **(A)** qRT-PCR analysis of relative gene expression of mitochondrial DNA (mtDloop and mtCO2) in the cytoplasmic fraction of siControl- and siGSDME-transfected SH-SY5Y cells post-CHPV infection. Nuclear GAPDH was used as an internal control. **(B)** Immunoblot analysis of total and phospho-STING protein levels in the whole-cell extracts of siControl and siGSDME-transfected SH-SY5Y cells infected with CHPV and analyzed at 24 hpi. (M=Mock and I=Infected). **(C)** Immunoblot analysis of total and cleaved GSDME-N terminal protein levels in the whole-cell extracts of siControl and siSTING-transfected SH-SY5Y cells infected with CHPV and analyzed at 24 hpi. (M=Mock and I=Infected). **(D)** Proposed model depicting the temporal regulation of mitochondrial stress, GSDME-localization to mitochondria, release of mtDNA and STING-dependent inflammatory signaling during CHPV infection.

Together, our findings support a temporally coordinated model linking mitochondrial stress, innate immune sensing, and inflammatory cell death during CHPV infection in neuronal cells (**Fig. 8D**). During the early phase of infection, MAVS depletion and attenuation of canonical RNA-sensing pathways coincide with limited downstream antiviral signaling. As infection progresses, mitochondrial dysfunction, characterized by increased mtROS and loss of mitochondrial membrane potential, is accompanied by caspase-3 activation and GSDME cleavage. The resulting GSDME-N fragment accumulates in mitochondria and contributes to mitochondrial membrane destabilization and mtDNA release, thereby promoting activation of the STING pathway. Consistent with this model, GSDME depletion reduced cytosolic mtDNA accumulation, STING phosphorylation, and CHPV-induced cell death, supporting a key role for GSDME in coupling mitochondrial perturbation to downstream inflammatory signaling. Notably, STING depletion also reduced GSDME abundance and cleavage, suggesting a possible reciprocal regulation between the GSDME and STING pathways.

## Discussion

The mechanistic understanding of viral encephalitis has been shaped predominantly by studies of neurotropic viruses belonging to the *Flaviviridae* family, including JEV, WNV and Zika virus (ZIKV) [64, 65]. These studies provided insights into viral neuroinvasion, neuroinflammation and neuronal injury. In contrast, considerably less is known about how members of the *Rhabdoviridae* family establish acute infection in the nervous system and, in particular, how they manipulate intrinsic neuronal immune pathways to promote viral replication and neuropathogenesis. CHPV, a neurotropic arbovirus, belonging to the *Rhabdoviridae* family, has emerged as a significant public health concern due to recurrent outbreaks in recent years with an exceptionally high fatality rate [66]. The virus is highly neuroinvasive, affecting children below fifteen years of age and causing rapid disease progression from an influenza-like illness to high-grade fever, seizures, coma and death within a narrow 24-48 hour window following the onset of symptoms [67]. Despite the severity of disease and recurrent outbreaks, the molecular mechanisms underlying CHPV-associated encephalitis remain poorly understood. Given the rapid and aggressive course of CHPV infection and other neurotropic rhabdoviruses, defining the temporal dynamics of antiviral responses is particularly important, as the timing and magnitude of these responses may influence disease progression and determine potential windows for therapeutic intervention.

Unlike professional immune cells, neurons rely on cell-intrinsic innate immune pathways to recognize and respond to invading viruses, yet how these pathways are engaged, modulated or subverted during infection with neurotropic viruses remains incompletely understood. Studies of neurotropic Flaviviruses have established the importance of the RIG-I/MDA5-MAVS-interferon axis in controlling viral infection and limiting neuropathogenesis [68]. In WNV infection, disruption of MAVS-dependent signaling compromises viral control and is associated with increased viral burden and mortality [69, 70] while studies of ZIKV have demonstrated direct viral antagonism of the RLR-MAVS pathway [71]. Similarly, neuronal RIG-I and STING signaling have been implicated in the induction of antiviral and inflammatory responses during JEV infection [33]. These observations establish suppression of cell-intrinsic antiviral signaling, particularly the MAVS-dependent interferon response, as an important strategy employed by neurotropic Flaviviruses to facilitate infection within the nervous system but such studies are limited in Rhabdovirus-mediated encephalitis like CHPV.

In the present study, we examined the innate immune response to CHPV infection in both neuronal cell lines and mouse brain and identified a distinct temporal pattern of antiviral and inflammatory signaling. Our findings demonstrate that CHPV infection is associated with an early impairment of the MAVS-dependent antiviral response accompanied by limited induction of both type I and type III IFN responses. In contrast, a marked induction of IFN and inflammatory responses emerged at later stages of infection, particularly at 24 hpi, despite the substantial reduction of MAVS protein levels. This suggests that the neuronal immune response to CHPV infection is not uniformly suppressed throughout the course of infection but undergoes a temporal transition from an early state characterized by impaired MAVS-associated antiviral signaling to a later state of robust innate immune activation. Importantly, this temporal pattern was also reflected in the global transcriptional landscape, revealing progressive remodeling of host gene expression at 4, 8 and 24 hpi in CHPV-infected neuronal cell lines and also in the mouse brain model at different stages of infection. To determine how this response was distributed across individual CNS cell populations *in vivo*, we performed scRNA-Seq analysis of whole brains from control and CHPV-infected mice. Our analysis revealed marked cell-type-specific heterogeneity in the antiviral response. Microglia exhibited a robust interferon-stimulated gene signature accompanied by induction of inflammatory mediators, whereas antiviral responses varied substantially across astrocyte and the neuronal sub-populations. However, the lower sequencing depth in our study may have reduced transcript capture and sensitivity for detecting low-abundance transcripts and therefore some cell-state-specific transcriptional changes may not have been captured. Nevertheless, the dataset provided sufficient resolution to identify major cellular populations and infection-associated transcriptional changes.

The delayed induction of IFN responses, together with the functional TBK-1/IRF3/IRF7 signaling despite substantial depletion of MAVS protein levels in both the CHPV-infected neuronal cell line and mouse models emphasized involvement of alternate host immune pathways in CHPV-induced neuropathogenesis. This made us speculate that the aggressive nature of CHPV infection could be associated with an inflammation-induced cell death program where the virus manipulates the host immune response to induce cell death and facilitate subsequent viral dissemination within the CNS. One of the pathways that has been shown to be protective during Flavivirus-induced encephalitis is the STING pathway [33, 72, 73]. However, in contrast to other encephalitis virus infections, our data suggest that CHPV induces robust STING phosphorylation in both neuronal cell line and mouse brain. Furthermore, pharmacological inhibition of STING or siRNA-mediated knockdown, confirmed that STING plays an important role in CHPV-induced neuroinflammation and the induction of pyroptotic cell death.

The cGAS-STING pathway is activated upon recognition of cytosolic dsDNA by cGAS. Although mtDNA leakage into the cytosol has been shown to activate the cGAS–STING pathway [74], whether mitochondrial damage and subsequent mtDNA release contribute to virus-induced neuroinflammation remains poorly understood. This raises the possibility that mitochondrial perturbation may provide a link between CHPV infection and activation of the cGAS–STING pathway. We therefore investigated whether mitochondrial homeostasis is altered during CHPV infection. The delayed inflammatory response and STING-induced pyroptosis following CHPV infection was accompanied by pronounced mitochondrial dysfunction. Beginning at the later stages of infection and becoming prominent at 24 hpi, CHPV induced loss of mitochondrial membrane potential, increased mtROS production and marked alterations in mitochondrial morphology. In parallel, we observed increased accumulation of mtDNA and TFAM in the cytosol, indicating loss of mitochondrial integrity and suggesting that mitochondrial damage may contribute to the activation of the cytosolic DNA-sensing pathway during late infection.

Since GSDME has been implicated in mitochondrial membrane damage and mtDNA release, we next asked whether GSDME might link the mitochondrial injury observed during CHPV infection to subsequent STING activation. Our findings extend the emerging literature on GSDM-mediated inflammatory cell death by identifying, for the first time in a neurotropic Rhabdovirus infection, a functional interplay between mitochondrial GSDME and STING signaling. The accumulation of cleaved GSDME at mitochondria coincided with mtDNA release and subsequent activation of the STING pathway, suggesting that mitochondrial GSDME may provide a mechanistic link between mitochondrial injury and cytosolic DNA sensing during CHPV infection. Consistent with this possibility, GSDME depletion markedly reduced STING phosphorylation, indicating that GSDME contributes to robust activation of the STING pathway during CHPV infection. Conversely, STING depletion substantially reduced the levels of cleaved GSDME, revealing a reciprocal relationship between these two pathways. Functionally, silencing either GSDME or STING attenuated CHPV-induced neuronal cell death and inflammatory responses, although their effects on viral burden differed. GSDME depletion was associated with a reduction in viral load, whereas STING depletion did not significantly alter overall viral burden, suggesting that STING contributes predominantly to the inflammatory and cell-death response rather than directly regulating CHPV replication.

These findings also raise questions regarding the contribution of the canonical inflammasome pathway to CHPV-induced inflammatory cell death. Although caspase-1 activation was detected, we did not observe convincing GSDMD cleavage under the conditions examined, while GSDME cleavage was prominent and closely associated with the inflammatory and lytic phenotype of infected neurons. Thus, the observed neuronal pyroptotic response cannot be readily explained by canonical GSDMD-dependent pyroptosis. At the same time, we cannot exclude a contribution of GSDMD, as activation of caspase-1 in the absence of detectable GSDMD cleavage does not by itself establish that GSDMD is dispensable. Further studies using genetic depletion of GSDMD, together with characterization of its subcellular activation and membrane permeabilization, will be required to define its precise contribution to CHPV-induced neuronal injury.

An additional important finding of this study is the cell-type-specific nature of the host response to CHPV infection. The inflammatory and cell-death pathways identified in infected neuronal cells do not necessarily reflect the responses of all infected or bystander cells within the brain. The distinct transcriptional and inflammatory signatures observed across cell populations suggest that CHPV infection engages a complex, cellular-context-dependent antiviral response in the central nervous system. Defining how neuronal, glial and other brain-resident cell populations differentially regulate mitochondrial stress, cGAS-STING signaling and GSDM activation will be important for understanding how local cellular responses collectively shape neuroinflammation and disease severity.

In conclusion, our findings identify mitochondrial dysfunction and the GSDME-STING axis as an important link between innate immune activation and neuronal death during CHPV infection where STING is the primary driver for CHPV-induced pyroptotis. This is in contrast with Flavivirus-induced encephalitis where STING has been shown to play a protective role. Future studies should determine how mitochondrial GSDME regulates mtDNA release, establish the precise contribution of GSDMD and other GSDMs to CHPV-induced cell death and define how these pathways crosstalk with STING-mediated pyroptosis. Such studies may ultimately determine whether modulation of the mitochondrial GSDME-STING axis can limit pathological inflammation and neuronal injury during CHPV infection and potentially other viral encephalitis, offering a promising host-directed therapeutic candidate.

## Materials and Methods

### Animal ethics statement

The animal experiments were performed after approval from the Institutional Animal and Ethics Committee (IAEC) of the National Brain Research Centre (Approval no. NBRC/IAEC/2025/217). Postnatal day 10 (P10) BALB/c pups were used in the experiments. The pups were handled in accordance with good animal practices as defined by the Committee for Control and Supervision of Experiments on Animals (CCSEA), Government of India. Mice were housed with their mothers throughout the experiment for milk feeding under a 12-hour light/dark cycle.

### Virus propagation and plaque assay

CHPV (Strain No. 1653514) isolated from a human patient in Nagpur was a kind gift by Dhrubajyoti Chattopadhyay (Sister Nivedita University, Kolkata, India). The virus was propagated in the Vero E6 cell line. Plaque assay was used to determine the titer of the virus and the titer was found to be 2.5 × 10^7^ PFU/mL and the infection dose was standardized accordingly.

### Animal treatment

P10 BALB/c pups were randomly assigned to four groups: Control (C), Day 1 post-infection (D1), Day 2 post-infection (D2) and full symptom (FS). Mice of either sex were administered with 3000 PFU of CHPV in a 50 μL volume through the intraperitoneal (i.p) route. Control animals were injected with phosphate-buffered saline (PBS). CHPV-infected animals showed symptoms by 72 to 96 hours post-infection, including weight loss, limb paralysis, body seizures, and restriction in movement. Symptoms scoring was conducted once daily until terminal illness or death (**Table 1**). Three animals for each group were collected at respective time points. The brain and spinal cord were excised after repeated transcardial perfusion with chilled 1X PBS and stored at -80°C until further use. The brains for cryosections were collected after transcardial perfusion with PBS, followed by tissue fixation using 4% paraformaldehyde (PFA). Brains were cryosectioned to obtain 20 µm transverse sections using a Leica CM3050 S cryostat.

**Table 1:** Symptom scoring after CHPV infection in mice.

| Score | Description |
| --- | --- |
| 0 | Normal Locomotion, Abdomen above ground |
| 1 | Tip-toe movement with elevated abdomen |
| 2 | Piloerection, Limping, Tip-toe movement |
| 3 | Piloerection, Hindlimb paralysis, Lowered abdomen, Movement Restriction, Seizures (Terminal Illness) |

### Cell lines

Human neuroblastoma cell line SH-SY5Y was a kind gift from Sourish Ghosh (Indian Institute of Chemical Biology, Kolkata) and the mouse microglia N9 cell line was a kind gift from Jayasri Das Sarma (Indian Institute of Science Education and Research, Kolkata). SH-SY5Y cells were cultured in DMEM-F12(1:1) (1X); #11320-033; Gibco) supplemented with 10% FBS (#16000-044; Gibco), 1% Penicillin-Streptomycin solution (#15140122; Gibco), 100 μg/ml Gentamicin solution (#A010; Himedia), 2.5 μg/ml Amphotericin B (#A011, Himedia). N9 cells were cultured in RPMI 1640 (#11875-093; Gibco) supplemented with 10% FBS, 1% Penicillin-Streptomycin solution, 2.5 μg/ml Amphotericin B. All cell lines were maintained at 37°C and 5% CO_2_ and were routinely tested for Mycoplasma by qRT-PCR using mycoplasma specific primers (**Table S1**).

### Antibodies and chemicals

Rabbit polyclonal antibody against MAVS (#14341-1-AP) was purchased from Proteintech. Rabbit polyclonal antibodies for pSTING (#19781), STING (#13647), pTBK1(#5483), TBK1(#3504), pIRF3 (#4947), IRF3 (#4302), were purchased from Cell Signaling Technology Inc. (CST). Rabbit polyclonal antibody against Gasdermin E (#13075-1-AP) and STING (#66680-Ig) mouse monoclonal antibody were purchased from Proteintech and cleaved caspase 3 (#ab2302) was purchased from Abcam. Calreticulin (#A1066), RIG-I (#A0550) and GM130 (#A5344) rabbit polyclonal antibodies were purchased from Abclonal. CHPV antibody was a kind gift from Debasis Nayak, IISER Bhopal. HRP tagged anti-mouse (#ab97023) and anti-rabbit secondary antibodies (#ab97051) were purchased from Abcam, anti-mouse DyLight800 (#5257P) was purchased from Cell Signaling Technology Inc. and Alexa Fluor 680 anti-rabbit (#A10043; Thermo Fisher Scientific) was used. For immunofluorescence staining, secondary antibodies Alexa Fluor 594 anti-rabbit (#ab150080) and Alexa Fluor 488 anti-mouse (#ab150113) from Abcam were used. Autophagy inhibitors chloroquine (#C6628; Sigma-Aldrich), bafilomycin (#HY-100558; MedChemExpress) and STING inhibitor, STING-IN-2 (#HY-138682; MedChemExpress) were used in this study.

### Tissue culture infectious dose assay (TCID) assay

1 × 10⁴ Vero E6 cells were seeded in 96-well tissue culture plates (Corning Inc.) and infected with CHPV stock serially diluted to 10^-12 (in DMEM containing 2 FBS) in quadruplets. Incubation was then done for 48 h post which the infected supernatant was discarded. The cells were then stained with 1% w/v crystal violet (#28376; Sisco-Research laboratories Pvt. Ltd.) solution for 15 minutes. The stain was washed under running tap water and the viral titer was calculated using the Spearman-Karber algorithm.

### Bulk RNA sequencing (RNA-Seq)

Total RNA was extracted from the indicated samples and assessed for concentration and integrity prior to library preparation. RNA sequencing libraries were prepared from samples meeting the required quality criteria and sequenced on an Illumina sequencing platform. Raw sequencing reads were subjected to quality control and preprocessing to remove adapter sequences and low-quality reads, followed by alignment to the appropriate reference genome and generation of gene-level read counts. Differential gene expression analysis was performed between the indicated experimental groups using normalized read counts. Genes with an adjusted p-value (false discovery rate, FDR) < 0.05 and an absolute log₂ fold change (|log₂FC|) ≥ 1 were considered significantly differentially expressed. Upregulated and downregulated genes were subsequently subjected to Gene Ontology and pathway enrichment analyses to identify biological processes and molecular pathways associated with the transcriptional changes.

### Single-cell RNA sequencing (scRNA-Seq)

Single-cell RNA sequencing data from the control and infected (FS stage) mouse brains were analyzed to characterize cell-type composition and gene expression patterns following CHPV infection. Annotated cell populations were visualized using UMAP plots with a manually assigned, consistent colour palette across datasets to facilitate comparison of cell-type distributions. The number and relative proportion of cells corresponding to each annotated cell type were calculated for each dataset and expressed as a percentage of the total cell population. To assess viral response- and immune-related gene expression, a curated panel of 106 genes was converted to mouse gene nomenclature and cross-referenced with the gene annotations of both Seurat objects. Genes without a corresponding annotation were excluded, resulting in a final panel of 93 genes common to both datasets. For differential gene expression analysis, the control and treated datasets were merged and cells were grouped according to the manual cell-type annotation. Differential expression between conditions was then assessed independently within each cell type using the Wilcoxon rank-sum test, with genes required to be expressed in at least 10% of cells and to meet a minimum log₂ fold-change threshold of 0.25. Cell types containing fewer than three cells in either condition were excluded from the analysis. Differentially expressed genes (DEGs) were ranked according to average log₂ fold-change and results were compiled separately for each cell type.

### Cell viability assay

Cell viability following CHPV infection was assessed using the 3-(4,5-dimethylthiazol-2-yl)-2,5-diphenyltetrazolium bromide (MTT) dye reduction assay (Sisco Research Laboratories Pvt. Ltd.). Briefly, 1 × 10⁴ cells were seeded in 96-well tissue culture plates (Corning Inc.) and infected with CHPV at the indicated multiplicity of infection (MOI). Where indicated, cells were pre-incubated for 24 h with chloroquine (50 µM), bafilomycin A1 (100 nM), or the STING inhibitor (0.5 µM and 1 µM) prior to CHPV infection. For gene-specific inhibition, cells were transfected with siRNA targeting STING (siSTING; 50 nM) or GSDME (siGSDME; 50 nM) for 24 h prior to infection with CHPV. At the indicated time points following infection, 10 µL of 5 mg/mL MTT solution was added to each well and cells were incubated for 4 h at 37°C. The medium was subsequently removed, and the resulting formazan crystals were dissolved in 150 µL DMSO by incubation for 15 min. Absorbance was measured at 570 nm using a microplate reader (BioTek, Agilent Technologies, USA). Each experiment was performed in triplicate, and background-corrected absorbance values were used to determine relative cell viability.

### Western blot analysis

2.0 × 10⁶ cells were seeded in 100 mm dishes for 24 h and infected with CHPV at an MOI of 0.1 for 1 h. Following removal of the virus, cells were harvested at the indicated time points and lysed in RIPA buffer supplemented with 1X protease and phosphatase inhibitors. Lysates were vortexed for 15 s at 5 min intervals for 30 min. Mouse brain tissues were collected in RIPA buffer supplemented with 1X protease and phosphatase inhibitors and homogenized using a bead beater (#85600; Qiagen). Proteins were precipitated with 20% TCA and incubated on ice for 1 h with intermittent vortexing. Following centrifugation at 14,000 rpm for 5 min at 4°C, pellets were washed twice with 500 µL ice-cold acetone, briefly vortexed, and centrifuged as above. Pellets were air-dried for 5 min and resuspended in 4X Laemmli sample buffer. Protein concentration was estimated using BCA protein estimation kit (#2603100011730; Bangalore GeNei). Equal amounts of protein were mixed with 4× Laemmli buffer to a final 1X concentration and denatured at 95°C for 5 min. Proteins were separated by SDS-PAGE and transferred onto PVDF membranes (Bio-Rad), followed by blocking with 5% skimmed milk or 5% BSA in 1X TBS. Membranes were incubated overnight at 4°C with appropriate primary antibodies, washed with 1X TBST, and incubated with the corresponding HRP or infrared/DyLight-conjugated secondary antibodies (Thermo Fisher Scientific) for 1 h at room temperature. Protein bands were detected using either ECL substrate (#170-5061, Bio-Rad) with a ChemiDoc MP Imaging System (Bio-Rad) or fluorescent detection using the LI-COR Odyssey DLx Imaging System. Band intensities were quantified using Image J software.

### Immunofluorescence and confocal microscopy

For immunofluorescence staining, 12-mm coverslips were coated with Geltrex™ (#A15696-01; Thermo Fisher Scientific) for 1 h at 37°C, followed by washing with 1× PBS. Cells (0.04 × 10⁶) were seeded onto coated coverslips and cultured for 24 h before treatment. Following treatment, cells were washed with 1× PBS and fixed with 4% paraformaldehyde for 12 min at room temperature. Cells were subsequently washed with 1× PBS and permeabilized with 0.2% Triton X-100 for 10 min at 4°C. Cells were blocked for 30 min at room temperature in blocking buffer containing 5% FBS, 0.05% Triton X-100, and 0.2 M glycine, followed by overnight incubation at 4°C with the respective primary antibodies. Cells were then incubated with appropriate Alexa Fluor 488- or Alexa Fluor 594-conjugated secondary antibodies and DAPI (Bio-Rad) for 1 h at room temperature. Coverslips were mounted using Fluoromount-G (#00-4958-02; Thermo Fisher Scientific) and imaged by confocal microscopy. For live-cell imaging, 1 × 10⁶ cells were seeded in 35-mm glass-bottom dishes and subjected to the indicated treatments. Cells were subsequently incubated with MitoSOX™ (#M36008; Invitrogen/Molecular Probes), TMRM, or Hoechst 33342 (Sigma-Aldrich) for 30 min in fresh culture medium, as appropriate, and immediately imaged by confocal microscopy.

For immunofluorescence analysis of mouse brain cryosections, sections were subjected to antigen retrieval in 1X citrate buffer ([catalogue number]) at 80–90°C for 20 min, followed by permeabilization with 0.5% Triton X-100 in PBS (PBXT) for 20 min. Sections were washed three times with PBS for 5 min each and blocked for 1 h at room temperature using 5% BSA and 5% FBS in 0.01% PBXT. Sections were incubated overnight at 4°C with the respective primary antibodies diluted in the blocking buffer. Following three washes with 0.1% PBXT for 5 min each, sections were incubated with Alexa Fluor 488- or Alexa Fluor 594-conjugated secondary antibodies together with DAPI (Bio-Rad) for 1 h at room temperature. Sections were subsequently washed three times with 0.1% PBXT and mounted using Fluoromount-G (#00-4958-02; Thermo Fisher Scientific), followed by coverslip placement and sealing. Images were acquired using a Leica DMi8 inverted confocal laser-scanning microscope (Leica Microsystems) equipped with 405-, 488-, and 594-nm laser lines and a 60× oil-immersion objective using Leica Application Suite (LAS) software. Where required, multiple optical z-stacks were acquired using the automated scanning mode and maximum-intensity projections were generated. Images were acquired at room temperature. Images were analyzed using ImageJ v1.8.0_172 (64-bit), and representative images were processed at 300 dpi using Adobe Photoshop v24.2.0.

### Immunohistochemical (IHC) analysis of mouse brain tissue

Mouse brain sections were stained with Harris hematoxylin and eosin (H&E) for histopathological analysis. Sections were deparaffinized with two changes of xylene for 10 min each and rehydrated sequentially in 100%, 95%, and 75% ethanol for 7, 2, and 2 min, respectively, followed by a brief rinse in distilled water. Sections were stained with Harris hematoxylin for 10 min and rinsed under running tap water for 10 min. Following differentiation by dipping in 90% ethanol, sections were counterstained with Eosin Y for 30-60 s. Sections were subsequently dehydrated through 90% and 100% ethanol for 5 min each, air-dried, and mounted using DPX mounting medium. Stained sections were examined by bright-field microscopy.

### Real-time quantitative PCR (qRT-PCR)

Total RNA was extracted from cultured cells using PureZOL™ RNA Isolation Reagent (#7326880, Bio-Rad) according to the manufacturer’s instructions. For mouse tissues, total RNA was extracted from brain and spinal cord samples following homogenization in PureZOL using a bead beater, followed by RNA isolation according to the same protocol. cDNA was synthesized from equal amounts of total RNA using iScript™ Reverse Transcription Supermix for RT-qPCR (#1708841, Bio-Rad), according to the manufacturer’s instructions. Quantitative real-time PCR (qRT-PCR) was performed using iTaq™ Universal SYBR® Green Supermix (#1725124, Bio-Rad). Primer sequences are provided in **Table S1**. β2-microglobulin (B2M) and β-actin (ACTB) were used as reference genes, and relative gene expression was calculated using the 2^−ΔΔCt method.

### Enzyme-linked immunosorbent assay (ELISA)

The concentrations of IL1β, IL6 and TNFα in culture supernatants from CHPV-infected SH-SY5Y cells were quantified using commercially available ELISA kits (IL-1β, #E-EL-H0149; IL-6, #E-EL-H6156; TNF-α, #E-EL-H0109; Elabscience), according to the manufacturer’s instructions. Culture supernatants from SH-SY5Y cells were collected at the indicated time points following CHPV infection and processed for cytokine estimation. Standard curves were generated using the standards provided with each kit, and cytokine concentrations in the samples were calculated by interpolation from the corresponding standard curves.

### Lactate dehydrogenase (LDH) assay

Lactate dehydrogenase (LDH) release into the culture supernatant was measured as an indicator of plasma membrane damage and cell lysis using a commercially available LDH cytotoxicity assay kit (#11644793001; Merck Roche). Briefly, culture supernatants were collected from control and CHPV-infected SH-SY5Y cells at the indicated time points and processed according to the manufacturer’s instructions. Appropriate controls, including untreated cells and maximum LDH release controls, were included for normalization. The enzymatic reaction was allowed to develop as specified by the manufacturer, and absorbance was measured at 490 nm using a multimode microplate reader (BioTek, Agilent Technologies, USA). LDH release was expressed as a percentage of maximum LDH release after background correction.

### siRNA transfection

siRNA for STING (#HY-RS13916), and Gasdermin E (#HY-RS05858) was purchased from MedChemExpress. As per the experimental requirement, 0.01 x 10^6^, 0.03 x 10^6^ or 0.1 x 10^6^ cells were seeded in a 96-well plate, 24-well plate or 6-well cell culture dishes (Nunc), respectively. Transfection was done with 50nM siRNA using Lipofectamine RNAiMAX (#13778075; Thermo Fisher Scientific) according to the manufacturer’s instructions. After 24 h of transfection cells were replenished with fresh growth medium, following which cells were infected with CHPV at an MOI of 0.1. After 1 h, the media was replaced and experiments were performed at the indicated time points.

### Subcellular fractionation of mitochondria

Mitochondrial and cytosolic fractions were prepared from SH-SY5Y cells using a mitochondrial fractionation kit (MedChemExpress) according to the manufacturer’s instructions. Briefly, 2.2 × 10⁶ cells were seeded in 100-mm culture dishes 24 h prior to infection. Following infection, cells were harvested and subjected to differential centrifugation according to the kit protocol to obtain mitochondrial and cytosolic fractions. The isolated fractions were collected and protein concentrations were determined using BCA protein estimation kit.

### Quantification of cytoplasmic mitochondrial DNA (mtDNA)

To quantify cytosolic mtDNA, 2 × 10⁶ cells were seeded in 100 mm dishes (Nunc) for 24 h and infected with CHPV. Following infection, 1 × 10⁷ cells were harvested, pelleted, and washed once with 1 ml ice-cold 1× PBS. The cell pellet was resuspended in 300 µL digitonin lysis buffer containing 150 mM NaCl, 50 mM HEPES (pH 7.4), 25 µg/mL digitonin, and protease and phosphatase inhibitors, and incubated at 4°C with rotation for 10 min. Lysates were centrifuged at 2,000 × g for 10 min at 4°C, and the supernatant was transferred to a fresh 1.5 ml microcentrifuge tube and centrifuged again at 2,000 × g for 20 min at 4°C. This clarification step was repeated until no visible pellet remained, and the final supernatant was collected as the cytosolic fraction. The initial pellet obtained following digitonin lysis was washed with ice-cold 1× PBS and centrifuged at 20,000 × g for 5 min at 4°C. The resulting pellet was resuspended in 300 µl NP-40 lysis buffer containing 150 mM NaCl, 50 mM HEPES (pH 7.4), 1% NP-40 supplemented with 1x protease and phosphatase inhibitors and incubated on ice for 30 min to obtain a crude mitochondrial/nuclear fraction. DNA was extracted from both fractions using the DNeasy Blood & Tissue Kit (#69504; Qiagen) and the abundance of mtDNA in the cytosolic fraction was quantified by qRT-PCR using mitochondrial-specific primers, mtDloop and mtCyclic oxygenase2 (mtCO2) (**Table S1**).

### Fluorescence-activated cell sorting (FACS) analysis

For flow cytometric analysis, 0.3 × 10⁶ cells were seeded in 6-well plates (Nunc) and cultured for 24 h before infection. Cells were infected with CHPV for 1 h, followed by removal of the virus and incubation in complete growth medium for the indicated time points. For TMRM and MitoSOX staining, cells were harvested by trypsinization and collected by centrifugation. Cell pellets were washed twice with 1× PBS and resuspended in 1× PBS containing TMRM (50 nM) or MitoSOX (5 µM). Cells were incubated for 30 min at 37°C in the dark, followed by two washes with 1× PBS. Cells were finally resuspended in 1× PBS and analyzed by flow cytometry.

### SEM imaging

SH-SY5Y cells were seeded on Geltrex-coated glass coverslips and cultured for the indicated period before transfection with siSTING (50 nM). Following 24 h of siRNA transfection, cells were infected with CHPV for the indicated time points. Cells were fixed with 2.5% glutaraldehyde prepared in 1× PBS for 1 h at 4°C and washed twice with PBS. Samples were subsequently post-fixed with 1% (v/v) aqueous osmium tetroxide for 20 min at room temperature, followed by washing with PBS and distilled water. Samples were dehydrated by sequential incubation in 30%, 50%, 60%, 80% and 100% ethanol for 5 min each and dried overnight in a vacuum desiccator. Samples were examined using a Zeiss Supra 55VP scanning electron microscope operated at 5.2 kV.

### Statistical analysis

All statistical analysis were carried out using GraphPad Prism version 9.5.1 for Windows (GraphPad Software, La Jolla, CA, USA). The results are presented as mean ± SEM of atleast three independent experiments or as indicated. Statistical significance was measured by one-way analysis of variance (ANOVA) with Tukey post-test. Asterisks indicate level of significance (*P < 0.05, **P < 0.01 and ***P < 0.001).

## Acknowledgements

We sincerely thank Sourish Ghosh (CSIR-Indian Institute of Chemical Biology, Kolkata, India), Debasis Nayak (IISER Bhopal, India) and Rubia Mondal (Institute of Health Sciences, Presidency University, Kolkata, India) for providing reagents. The authors thank Rupak Datta (IISER Kolkata) and IISER Kolkata central facility staff member Mr. Kashinath Sahu for SEM imaging. The work was funded by extramural funding from Council for Scientific and Industrial Research (CSIR), Govt. of India (#37WS (0001) /2023-24/EMR-II/ASPIRE dated 20/6/2024) to PM. Research in AB’s lab is supported by the J.C. Bose Fellowship (JCB/2020/000037), Anusandhan National Research Foundation (ANRF), Government of India.

## Author contribution

PM and AS conceived the work and NK, SD, AP, JB, SR, SS, SC and JR performed the experiments. AP and AB assisted with designing and conducting the animal experiments. JR and DM assisted with the pyroptotic assays. SS assisted with the SEM experiments. NK, SD, AP, AS and PM wrote the manuscript and prepared the figures and AS and PM revised the entire manuscript.

## Declaration of AI in the writing process

During the preparation of this work, the author(s) used ChatGPT (https://chatgpt.com/) to improve the English or organize the results section, but not to draw any scientific conclusions.

## Supplementary figure legends

**Figure S1.**
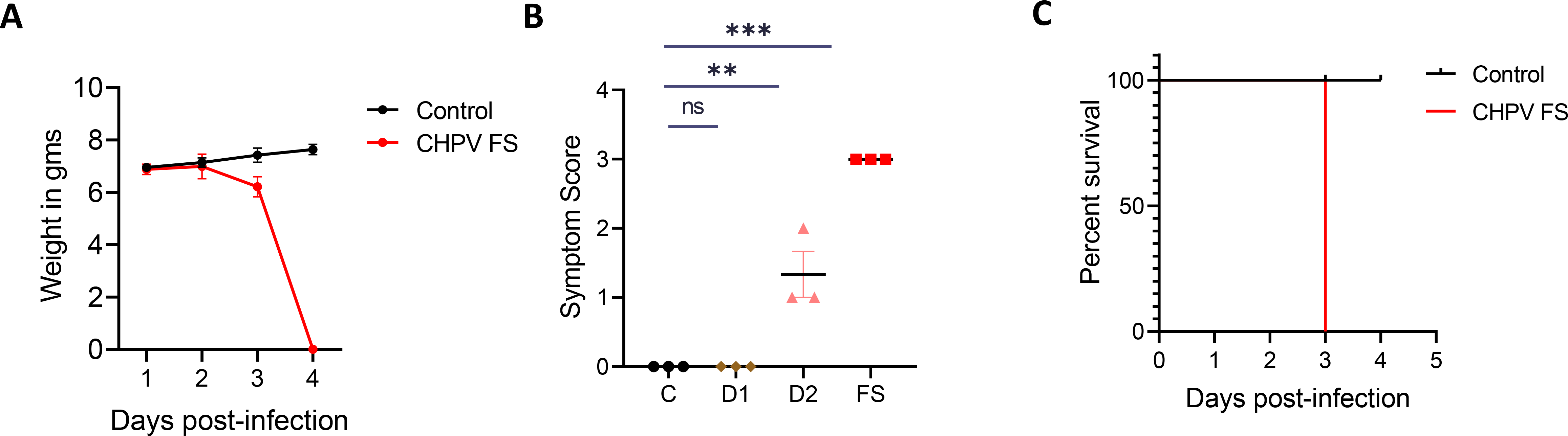
Disease progression in CHPV-infected postnatal day 10 (P10) BALB/c pups. **(A)** Body weight changes post-viral infection. CHPV-infected animals (red) show progressive decline in body weight as compared to control animals (black). **(B)** Survival proportion of Control (black) versus CHPV-infected group (red). All the CHPV-infected animals showed terminal illness on day 3 post-infection. **(C)** Symptom scoring across different groups shows progressive locomotion defect, with highest score observed at full symptom. Data are presented as mean ± SD. Statistical significance was determined by one-way ANOVA followed by Tukey’s multiple comparison test (*P < 0.12; **P < 0.01; ***P < 0.001; n=3 animals/group).

**Figure S2.**
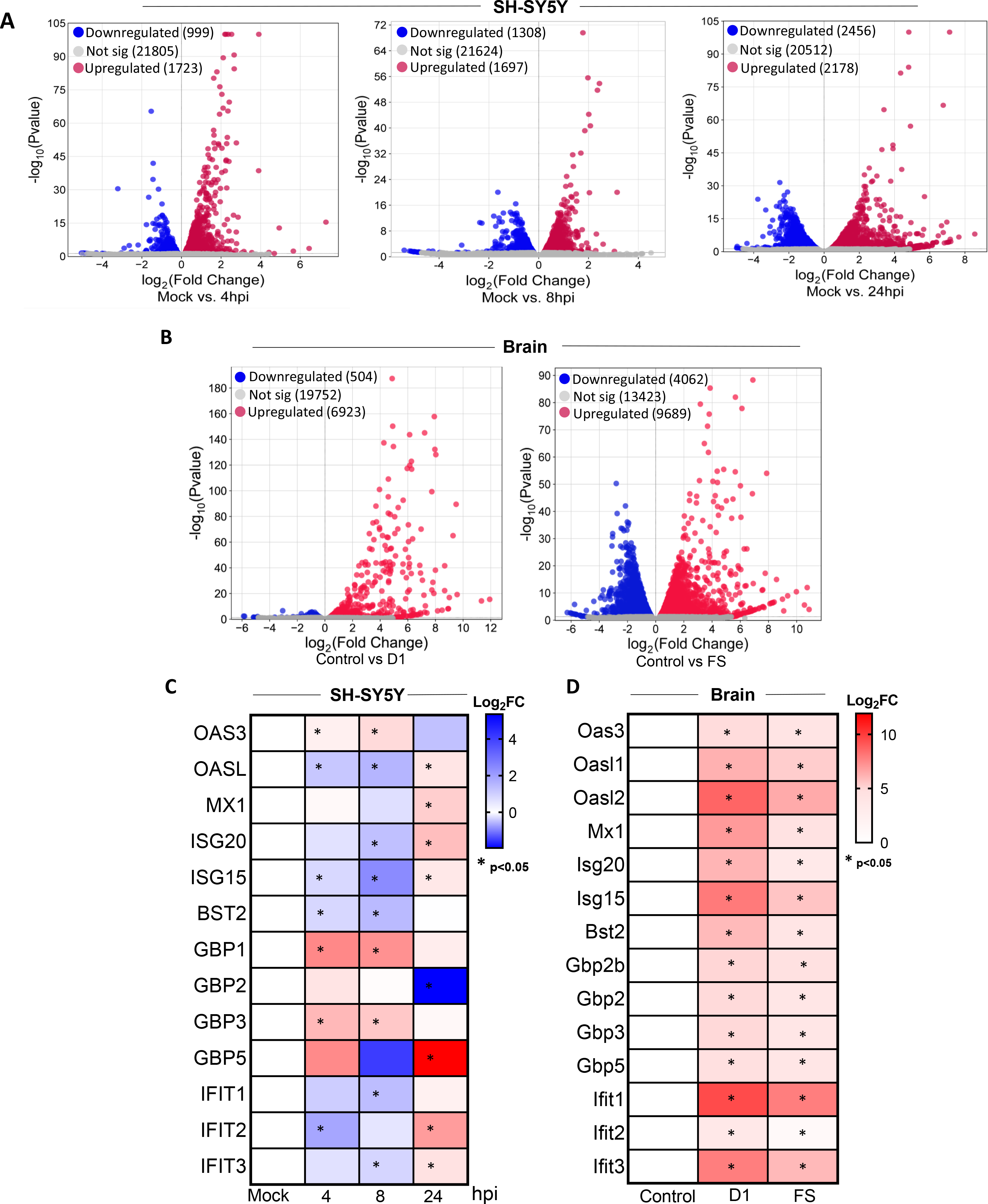
Differentially expressed genes from CHPV-infected SH-SY5Y cells and mouse brain. **(A)** Volcano plot showing mRNA expression of differentially expressed genes in the bulk RNA-Seq data from uninfected SH-SY5Y cells compared to CHPV-infected cells at 4 (left panel), 8 (middle panel) and 24 (right panel) hpi. Dotted line represents a threshold of 0 for log2FC. **(B)** Volcano plot showing differentially expressed genes from bulk RNA-Seq data of CHPV-infected mouse brain at D1 (left panel) and FS (right panel). **(C)** Heatmap showing relative mRNA expression levels of differentially expressed genes of ISGs at indicated time points obtained from bulk RNA sequencing in CHPV-infected SH-SY5Y cells. **(D)** Heatmap showing relative mRNA expression levels of differentially expressed genes of ISGs at indicated time points obtained from bulk RNA sequencing in CHPV-infected mouse brain tissue. * Represents the significant genes (p≤0.05).

**Figure S3.**
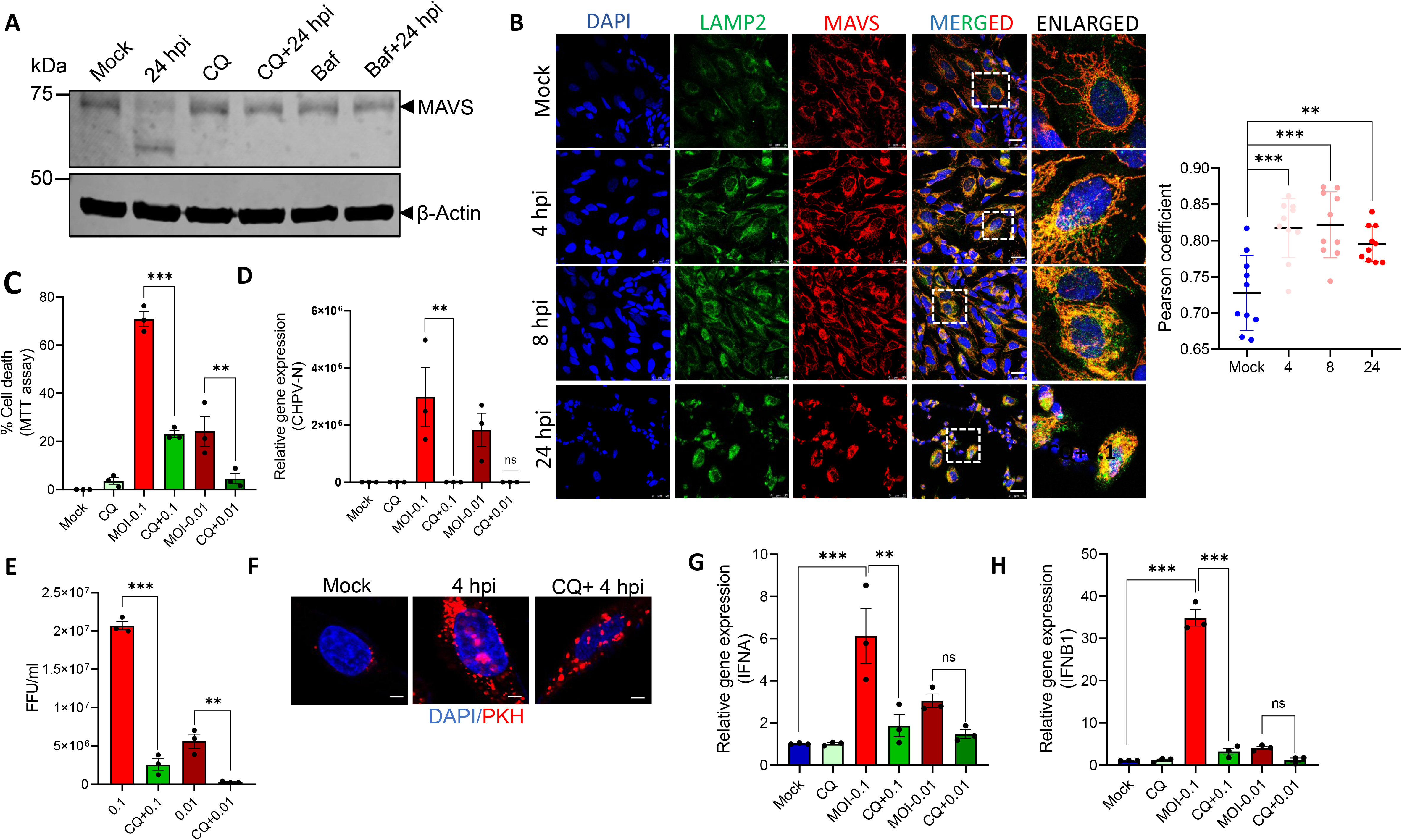
Autophagy inhibitors stabilizes MAVS at the mitochondria and prevent CHPV-induced cell death. **(A)** Immunoblot analysis of MAVS protein levels in the whole-cell extracts of CHPV-infected SH-SY5Y cells at 24 hpi in the presence or absence of the autophagy inhibitors Chloroquine and Bafilomycin. β-actin was used as loading control. **(B)** (Left panel) Representative confocal images showing lysosomal marker LAMP2 (green), MAVS (red) and DAPI (blue) in CHPV-infected SH-SY5Y cells. (Scale bar = 25µm). (right panel) Pearson’s correlation coefficient (PCC) showing the co-localization of MAVS with LAMP1. (n = 100 cells). **(C)** MTT assay showing the percentage of cell death following infection with CHPV at MOI 0.1 or 0.01 in the presence or absence of Chloroquine. **(D)** qRT-PCR analysis of the relative gene expression of CHPV N gene post-CHPV infection (MOI-0.1 and 0.01) at 24 hpi compared with mock-infected cells in the presence or absence of CQ. **(E)** Bar graph shows viral titre following infection with CHPV in the presence or absence of CQ. **(F)** Representative confocal image showing virus stained with the PKH dye (red) at 4 hpi in the presence or absence of CQ or Baf. (Scale bar = 5 µm). **(G-H)** qRT-PCR analysis of the relative gene expression of IFNA (G)and IFNB (H) genes post-CHPV infection (MOI-0.1 and 0.01) at 24 hpi compared with mock-infected cells in the presence or absence of Chloroquine. Data are presented as mean ± SD. Statistical significance was determined by one-way ANOVA with post hoc Tukey’s multiple comparison test (**P < 0.01; ***P < 0.001; n=3 biological replicates).

**Figure S4.**
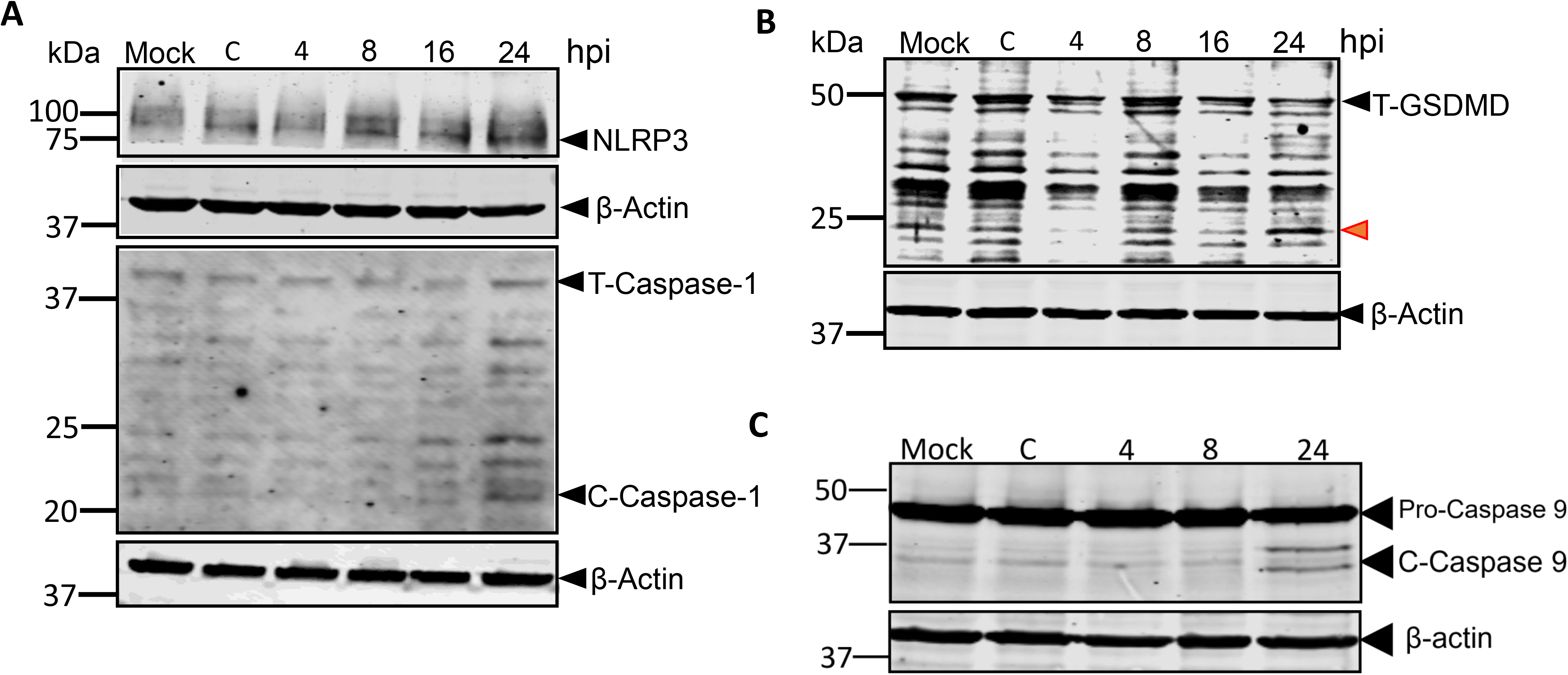
CHPV-infection associated NLRP3 inflammasome in SH-SY5Y cells. **(A)** Representative immunoblot images showing NLRP3, total and cleaved Caspase-1 levels following CHPV infection at 4, 8, 16 and 24 hpi. β-actin was used as loading control. **(B)** Immunoblot analysis of GSDMD and GSDMD-N at 4, 8, 16 and 24 hpi with CHPV. β-actin was used as loading control. **(C)** Immunoblot analysis of caspase 9 (total and cleaved), in the whole-cell extracts of CHPV-infected SH-SY5Y cells at 4, 8 and 24 hpi and compared with mock-infected and control (C) cells. β-Actin and GAPDH was used as a loading control. Data are presented as mean ± SD. Statistical significance was determined by one-way ANOVA with post hoc Tukey’s multiple comparison test (**P < 0.01; ***P < 0.001; n=3 biological replicates).

**Figure S5.**
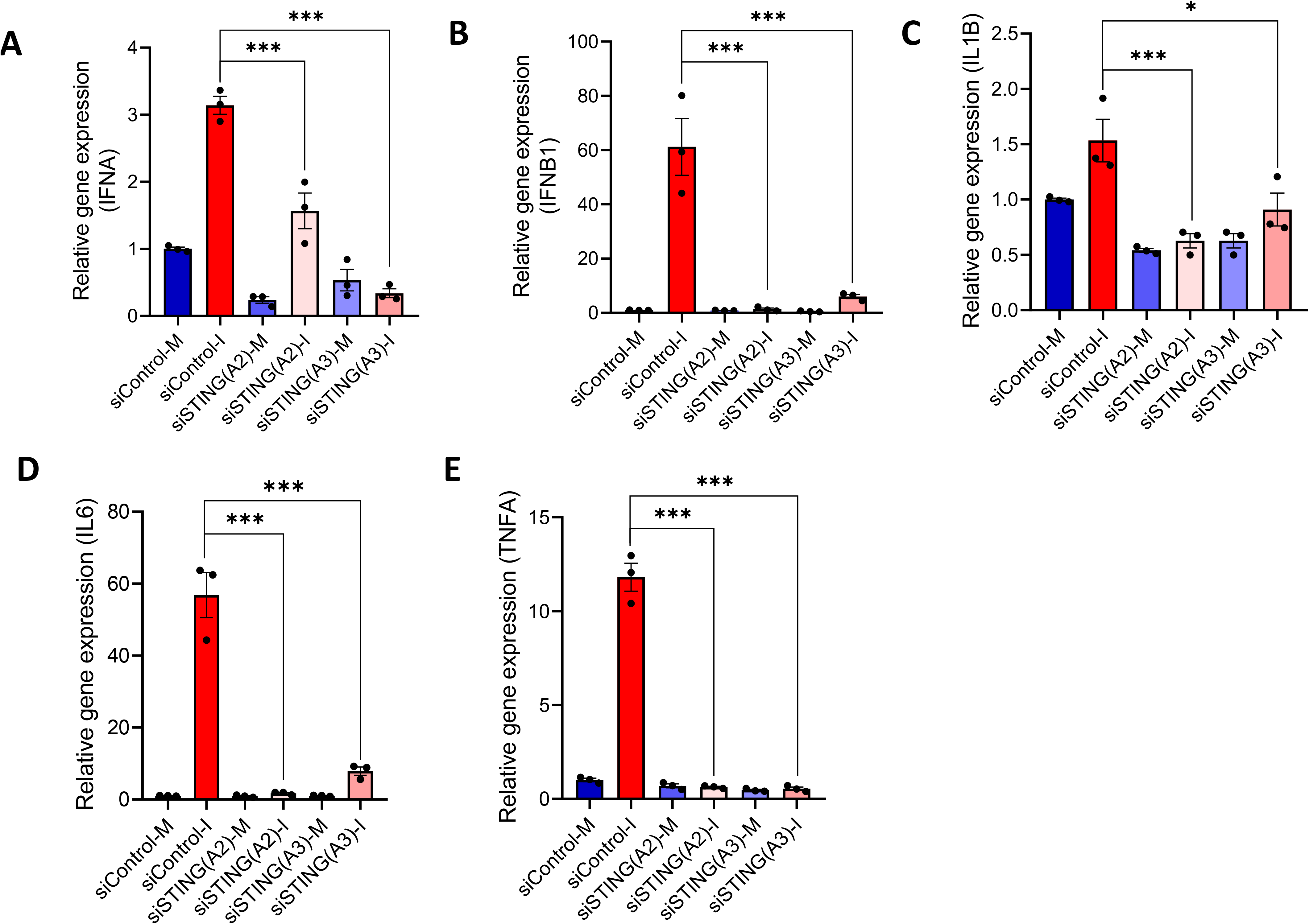
siSTING reduces induction of type-I interferons and inflammatory responses in CHPV-infected SH-SY5Y cells. **(A-B)** qRT-PCR analysis of the relative gene expression of IFNA (A)and IFNB (B) genes post-CHPV infection (MOI-0.1) at 24 hpi in siControl and siSTING transfected samples. **(C-D)** qRT-PCR analysis of the relative gene expression of IL1B (C), TNFA (D) and IL6 (C) genes post-CHPV infection (MOI-0.1) at 24 hpi in siControl and siSTING transfected samples. (M = mock; I = infected). Data are presented as mean ± SD. Statistical significance was determined by one-way ANOVA with post hoc Tukey’s multiple comparison test (**P < 0.01; ***P < 0.001; n=3 biological replicates).

**Figure S6.**
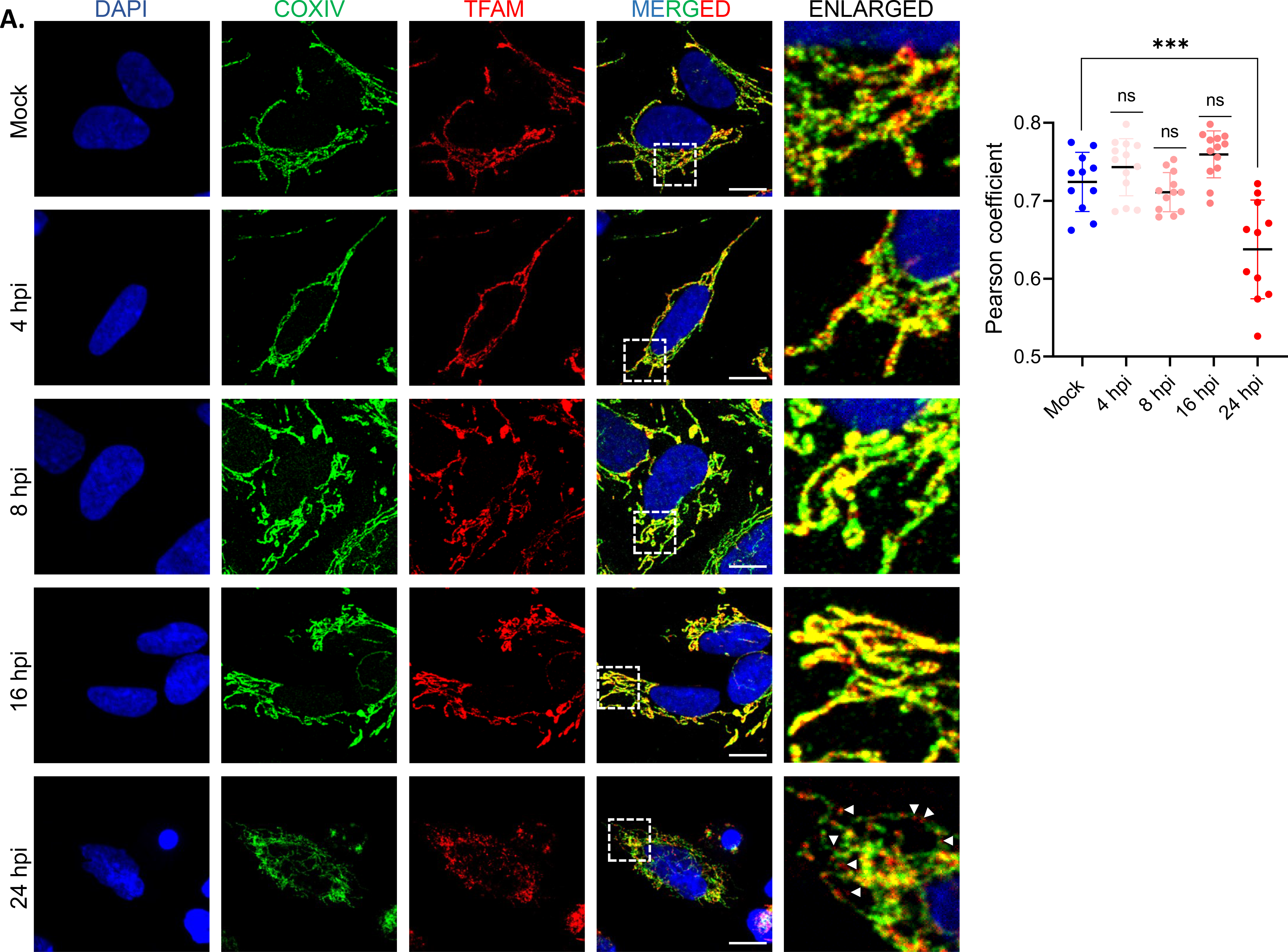
(Left Panel) Representative immunofluorescence image showing mitochondrial membrane marker COXIV (green) and mitochondrial DNA marker TFAM (red) in CHPV-infected SH-SY5Y cells at the indicated times. (Scale bar= 10 µm). (Right panel) Colocalization of COXIV and TFAM is shown through Pearson’s correlation coefficient (PCC) (n=100 cells). Data are represented as Mean ± SD. Statistical significance was determined by one-way ANOVA with Tukey’s multiple comparison test (***P < 0.001)

**Figure S7.**
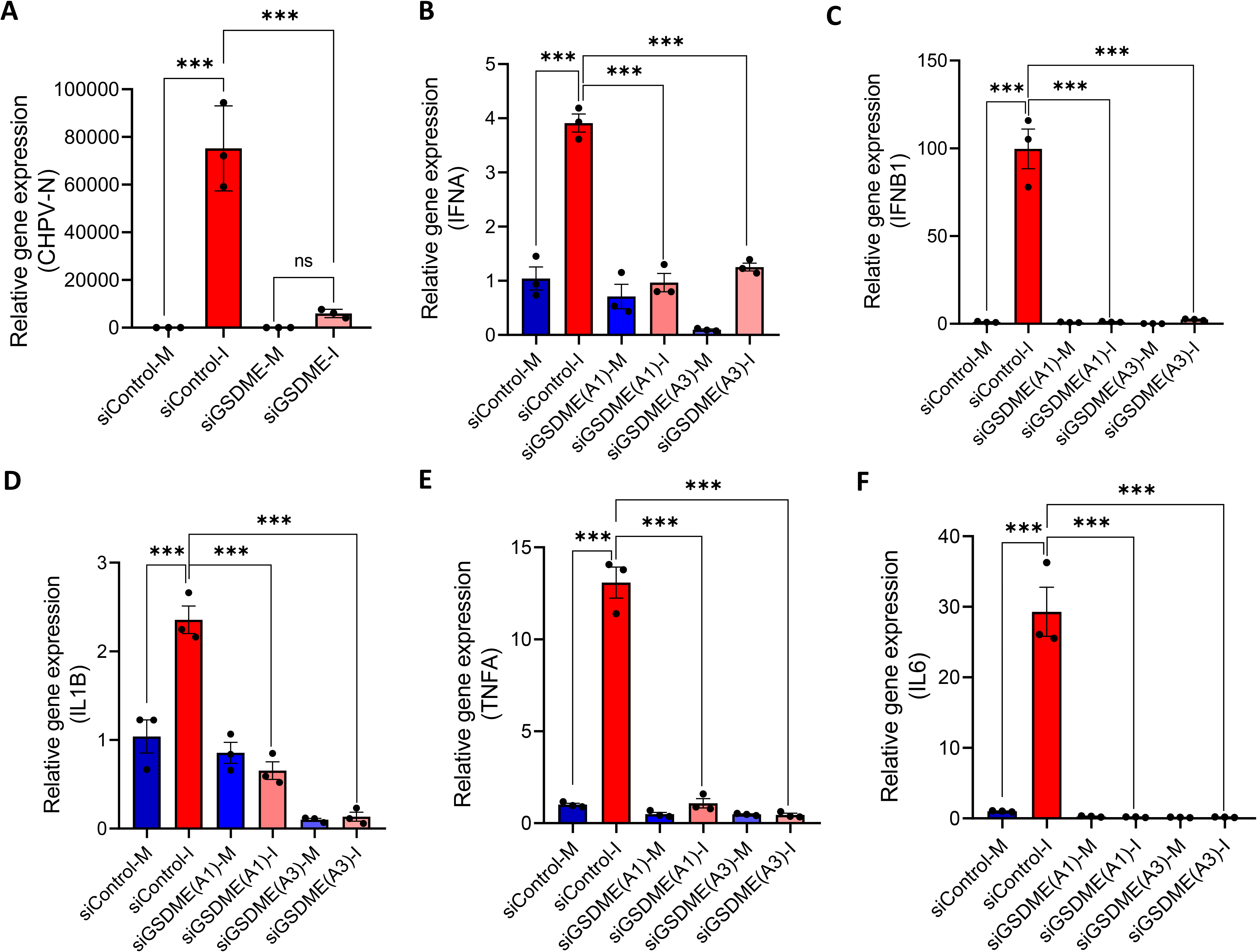
**(A)** qRT-PCR analysis of the relative gene expression of CHPV N gene in SH-SY5Y cells transfected with siGSDME for 48 h and analyzed at 24 hpi. **(B)** qRT-PCR analysis of the relative gene expression of IFNA and IFNB in SH-SY5Y cells transfected with siGSDME for 48 h and analyzed at 24 hpi. **(C)** qRT-PCR analysis of the relative gene expression of IL1β, TNFα, and IL6 in SH-SY5Y cells transfected with siGSDME for 48 h and analyzed at 24 hpi. (M = mock; I = infected)

